# A compendium of adipocyte morphologies across different breast pathologies

**DOI:** 10.1101/2025.08.08.669287

**Authors:** Abigail Dodson, Katie Hanna, Kerri Palmer, Hafeez Ibrahim, Rasha Abu-Eid, Gerald Lip, Nicola Spence, Ehab Husain, Beatrix Elsberger, Justin J. Rochford, Valerie Speirs

**Affiliations:** Institute of Medical Sciences, School of Medicine, Medical Sciences and Nutrition, University of Aberdeen, Aberdeen, AB25 2ZD, UK; Institute of Dentistry, School of Medicine, Medical Sciences and Nutrition, University of Aberdeen, Aberdeen, AB25 2ZD, UK; The Rowett Institute, School of Medicine, Medical Sciences and Nutrition, University of Aberdeen, Aberdeen, AB25 2ZD, UK; Aberdeen Royal Infirmary, Aberdeen, AB25 2ZN, UK; Raigmore Hospital, Inverness, IV2 3UJ

**Keywords:** Adipose tissue biology, adipose remodelling, lipolysis, fatty acids, biomarkers

## Abstract

Adipocytes are abundant in the breast tissue microenvironment. In breast cancer, they can change morphologically according to their proximity to tumour cells, with the closest becoming cancer-associated adipocytes (CAAs). It remains unclear whether breast cancer risk factors, including menopausal status, body mass index (BMI), and mammographic density (MD), influence CAAs morphology in breast carcinogenesis. This study aimed to quantify morphological differences in adipocytes across breast cancer pathologies and associated risk factors.

Whole slide images of hematoxylin and eosin stained cancer (*n* = 149) and normal (*n* = 182) breast tissue samples were analysed. Parameters representative of adipocyte morphology: perimeter, area, concavity, and aspect ratio, were measured using ImageJ. Adipocytes were considered close (≤ 2 mm) or distant (>2 mm) to cancer cells in cancer samples or breast epithelial cells in normal samples. Close adipocytes in cancer samples were designated CAAs.

CAAs decreased in size compared to distant adipocytes (*p ≤* 0.0001). A similar trend was observed between close and distant adipocytes in normal (*p ≤* 0.0001). CAAs size increased post menopause (*p* ≤ 0.0001). CAAs size positively correlated with BMI (*p* ≤ 0.0001). In cancer cases, distant adipocyte size increased and concavity decreased with increasing MD (*p* ≤ 0.01). Smaller CAAs were associated with poorer survival (*p ≤* 0.05).

Morphological differences were identified in adipocytes dependent on location within the breast, tissue, pathology and risk factors. Understanding what drives these morphological differences could provide mechanistic insight into whether risk factor-induced alterations in adipocytes influence their role in breast carcinogenesis.

## Introduction

Adipose tissue represents a major cellular component of the breast and has important endocrine functions [1, 2]. It is comprised of adipocytes and a collagen-rich extracellular matrix, in which other cell types are embedded including macrophages, preadipocytes, and fibroblasts [3]. White adipocytes are the most common type of adipocyte within the breast adipose tissue [2]. These highly specialised cells store energy, principally as triacylglycerol, which can be broken down into glycerol and fatty acids to meet localised or systemic energy requirements [4]. In breast cancer, adipocytes close to the tumour leading edge are known as cancer-associated adipocytes (CAAs). Breast cancer cells can induce adipocyte lipolysis resulting in CAAs, which are smaller, more elongated and irregular in shape due to the reduction in lipid storage [5–8]. Previous findings have highlighted differences in adipocyte area and/or diameter between CAAs and distant adipocytes, which are typically anywhere between 200 µm - 2 cm further away from the tumour leading edge [9–12]. Studies have found that adipocyte size can vary between the different types of breast cancer[13, 14]. CAAs and distant adipocytes from patients with invasive lobular cancer were smaller than those from invasive ductal carcinoma (IDC) and distant adipocyte size in primary ductal carcinoma *in situ* was positively correlated with the risk of developing ipsilateral IDC [13, 14].

Breast cancer risk factors, including menopausal status, obesity, and mammographic density (MD) can influence breast adipose tissue [15–20]. However, the relationship between these three risk factors is complex. MD and BMI have an inverse relationship in terms of breast cancer risk [21–23]. As MD increases, the proportion of adipose tissue in the breast decreases, whereas an increase in BMI is associated with a rise in adipose tissue. Despite the differences in adipose tissue volume, both these factors increase the risk of breast cancer development. A high MD, defined as breast tissue of > 75% fibroglandular tissue, increases a female’s risk by 1- to 6-fold compared to a low MD, where breast tissue is mostly adipose (< 25% fibroglandular tissue) [24, 25]. The underlying mechanisms as to why higher MD poses an increased risk of developing breast cancer are yet to be fully elucidated. Moreover, studies have found that obese post-menopausal females have a higher risk of developing breast cancer compared with obese pre-menopausal females [26–28]. The reasons for this are unclear [29, 30].

During weight gain, adipose tissue expands its storage capacity to accommodate excess fat in the body through the generation of new adipocytes from precursor cells (hyperplasia) and hypertrophy of pre-existing adipocytes [15]. Several studies have revealed the association of obesity with adipocyte hypertrophy in the breast, where the diameter of adipocytes was positively correlated with both the severity of breast inflammation and Body Mass Index (BMI) [16–19]. The changes in adipocytes during obesity and the promotion of a pro-inflammatory state could provide a favourable environment for breast cancer development.

Post menopause, adipose tissue becomes the main source of estrogen as ovarian production ceases [31]. This is exacerbated with weight gain as an accumulation of adipose tissue within the breast can increase aromatase expression [32]. This enzyme catalyses the conversion of androgens to estrogens and increases the level of locally available estrogen by up to 10-fold higher than systemic levels [33–35]. These local changes in hormone levels have been reported to influence adipocyte morphology, with a stronger positive correlation between adipocyte diameter and aromatase expression in post-menopausal females compared with pre-menopausal females [20].

It is not fully understood whether menopausal status, BMI, and particularly MD influence the typical reduced size and less spherical shape of CAAs during breast cancer development. Therefore, this study used digital pathology to characterise morphological differences in adipocytes in relation to these three breast cancer risk factors. By including both cancer and normal samples, the study also examined how these risk factors impact adipocyte size with and without the influence of cancer, providing insight into whether these risk factors influence the contributions of altered adipocyte morphology to breast cancer risk, development, and progression.

## Results

### Cancer-associated adipocytes are morphologically distinct from close normal adipocytes

Adipocyte morphology was evaluated in normal and breast cancer tissue samples. CAAs showed a 28.7% decrease in size compared with distant adipocytes (*p* ≤ 0.0001; **Fig. 1a, b**). A similar trend in size was observed in adipocytes close to breast epithelial cells in normal samples (13.9%) (*p* ≤ 0.0001), as shown in H&E-stained cancer (**Fig. 1e, f, g**) and normal (**Fig. 1h, i, j**) breast tissue sample images. Close adipocytes also had lower concavity (*p* ≤ 0.0001; **Fig. 1c**) and higher aspect ratios (normal, *p* ≤ 0.05; cancer, *p* ≤ 0.0001; **Fig. 1d**) irrespective of the tissue pathology, meaning they were more elongated than distant adipocytes. CAAs were smaller (*p* ≤ 0.0001; **Fig. 1a, b**), less concave (*p* ≤ 0.0001; **Fig. 1c**), and more elongated (*p* ≤ 0.001; **Fig. 1d**) than close adipocytes from normal samples. The additional parameters discussed in the Random Forest model further support these findings as CAAs had a smaller convex area, feret, and breadth compared to normal close adipocytes (**Supplementary Fig. 1a, b, c**). Overall, these findings suggest that adipocytes have altered morphology in relation to their location within the breast and according to tissue pathology.

**Fig. 1.**
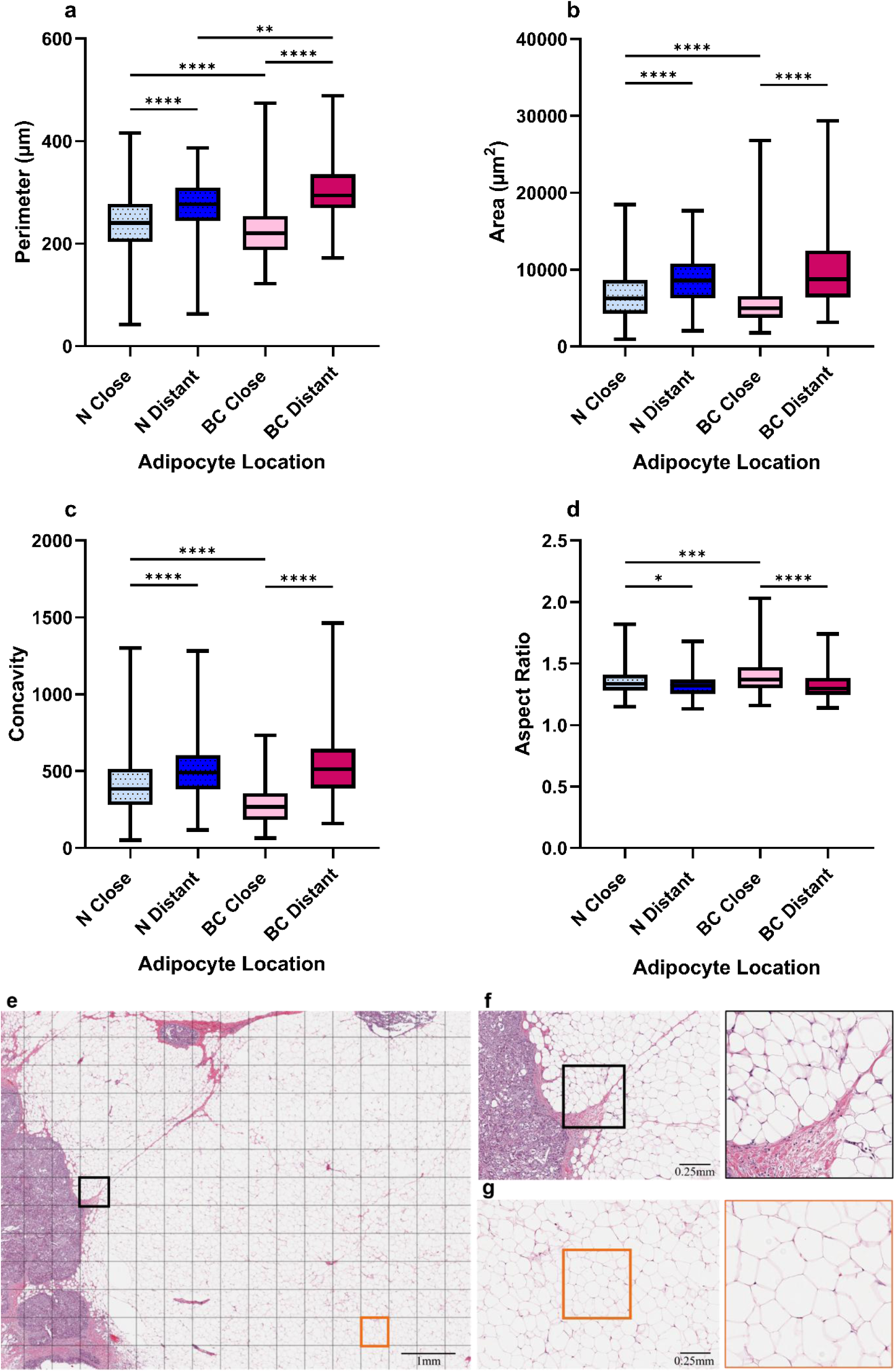
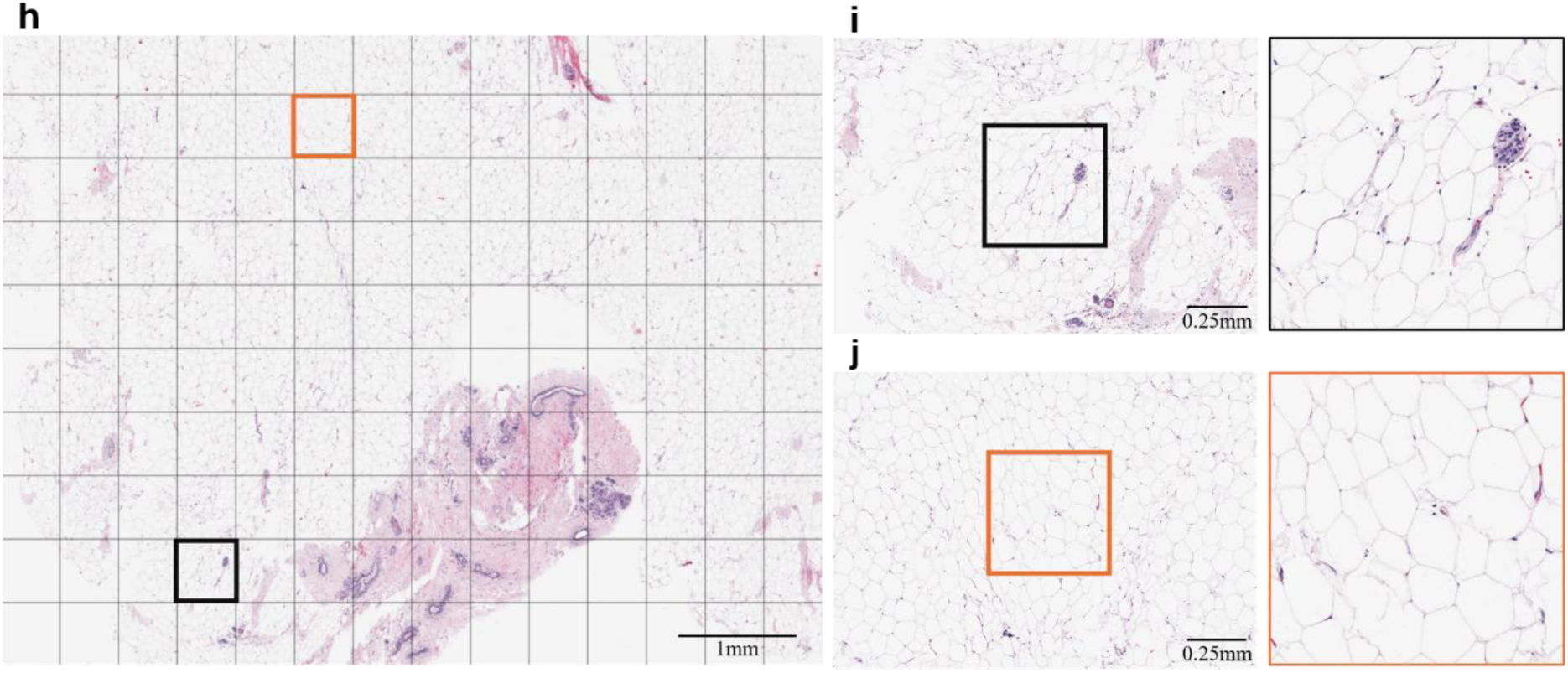
Characterisation of close and distant adipocyte morphology from BC and normal (N) breast tissue. Median adipocyte **a** perimeter, **b** area, **c** concavity, and **d** aspect ratio from breast cancer (*n* = 149) and normal (*n* = 153) samples. Representative examples of WSIs of H&E-stained **e** breast cancer and **h** normal tissue samples with **f**, **i** close (black squares) and **g, j** distant (orange squares) regions of interest (ROIs). N Close = normal adipocytes ≤ 2 mm away from the breast epithelial cells (*n* = 21,318). N Distant = normal adipocytes > 2 mm away from the breast epithelial cells (*n* = 11,274). BC Close = breast cancer adipocytes ≤ 2 mm away from the tumour-leading edge (*n* = 51,751). BC Distant = breast cancer adipocytes > 2 mm away from the tumour-leading edge (*n* = 27,003). * *p* ≤ 0.05, ** *p* ≤ 0.01, *** *p* ≤ 0.001, **** *p* ≤ 0.0001

### Adipocyte size differs according to menopausal status

Samples were stratified into pre- and post-menopausal groups to assess the potential impact of menopausal status on adipocyte morphology. Menopausal status was known for 97 cancer cases (16 pre- and 81 post-menopausal) and 130 normal cases (48 pre- and 82 post-menopausal).

In breast cancer tissue, close adipocytes were smaller and less concave than distant adipocytes (*p* ≤ 0.0001), irrespective of menopausal status (**Fig. 2a, b, c**). This trend was also observed in normal samples (pre, *p* ≤ 0.0001; post, perimeter and area = *p* ≤ 0.0001, concavity = *p* ≤ 0.001; **Supplementary Fig. 2a, b, c**). Comparison of size between menopausal groups in breast cancer samples showed that CAAs from pre-menopausal samples were smaller than CAAs and distant adipocytes from post-menopausal samples (*p* ≤ 0.05, and *p* ≤ 0.0001, respectively; **Fig. 2a**). No differences were seen between menopausal groups in normal samples. Limited differences in aspect ratio, except in breast cancer samples where post-menopausal distant adipocytes were less elongated than close adipocytes from pre-(*p* ≤ 0.001) and post-menopausal (*p* ≤ 0.0001) samples (**Fig. 2d**). In normal samples, distant adipocytes were less elongated than close adipocytes in only pre-menopausal samples (*p* ≤ 0.05; **Supplementary Fig. 2d**). These results indicate that cancer adipocyte morphology is influenced not only by their location within the breast and the tissue pathology but also by menopausal status, with CAAs being smaller in the pre-menopausal group compared with the post-menopausal group.

**Fig. 2.**
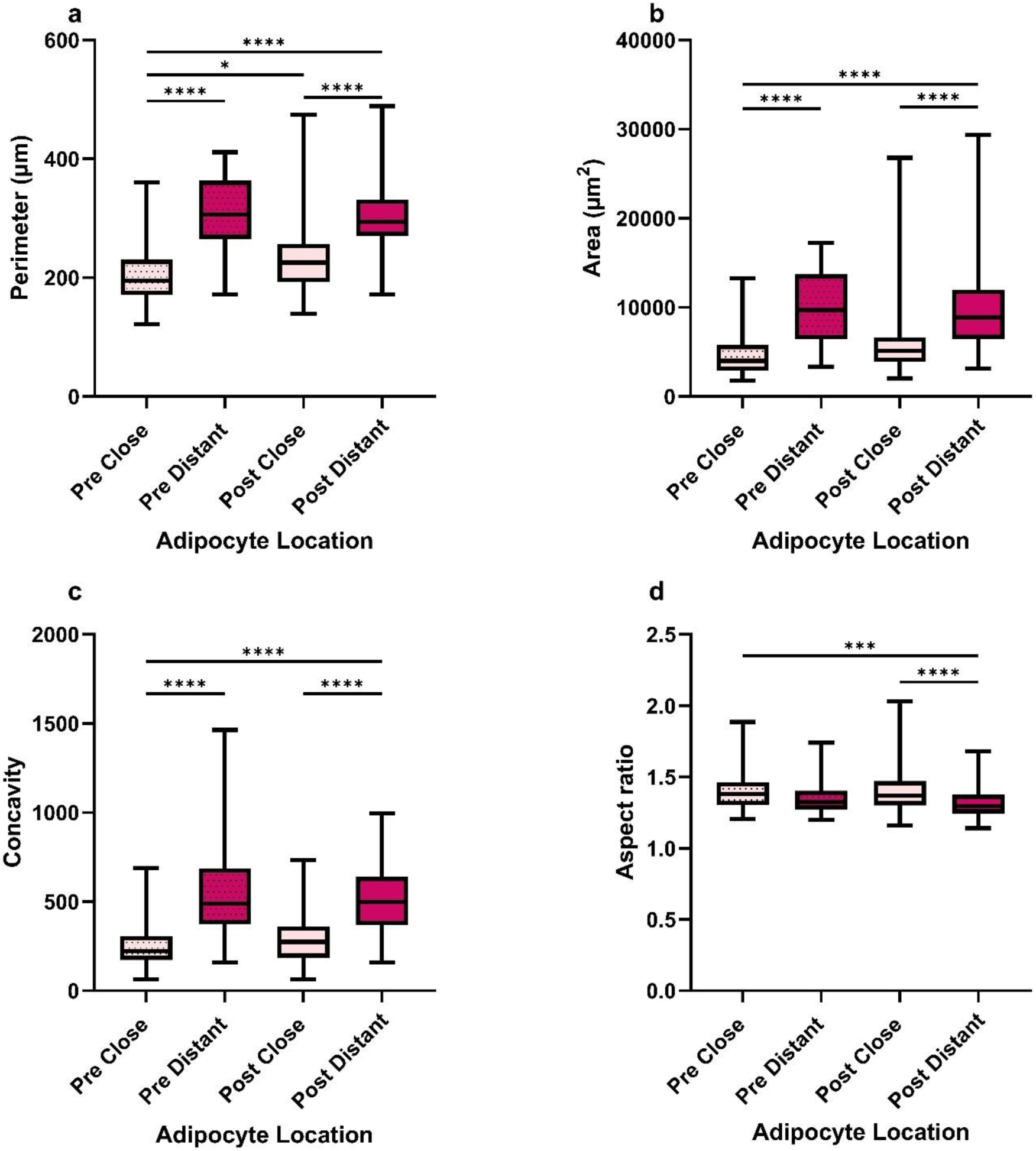
Pre-menopausal cancer-associated adipocytes were smaller than post-menopausal adipocytes. Median adipocyte **a** perimeter, **b** area, **c** concavity, and **d** aspect ratio from breast cancer tissue samples (*n* = 97). Pre = Pre-menopausal samples (*n* = 16). Post = Post-menopausal samples (*n* = 81). Close = Breast cancer adipocytes ≤ 2 mm away from the tumour-leading edge (pre, *n* = 2,868; post, *n* = 18,313). Distant = Breast cancer adipocytes > 2 mm away (Pre, *n* = 724; post, *n* = 3,135). * *p* ≤ 0.05, *** *p* ≤ 0.001, **** *p* ≤ 0.0001

### Adipocyte size is positively correlated with BMI

Adipocyte size increases with increasing BMI in normal breast tissue [17, 18]. Several studies have also noted this link between adipocyte hypertrophy and increased BMI in breast cancer [16–19]. However, it is not fully known how obesity may alter adipocyte shape during breast cancer. Therefore, we stratified 50 cancer and 84 normal samples into quartiles of BMI (group 1 < 18.5; group 2 18.5 - 24.9; group 3 25 - 29.9; and group 4 > 30). For cancer cases, 3 were in group 1, 17 in group 2, 10 in group 3, and 20 in group 4. For normal cases, 0 were in group 1, 27 in group 2, 26 in group 3, and 31 in group 4.

Close and distant adipocyte size increased from the low to high BMI group in cancer cases (perimeter, *p* ≤ 0.0001; area, *p* < 0.001; **Table 1; Supplementary Fig. 3a, b, c, d**). A similar trend was seen in normal close adipocyte size from group 2 and 4 (perimeter, *p* ≤ 0.01; area, *p* ≤ 0.0001; **Supplementary Fig. 4a, c**). Normal distant adipocytes from the highest BMI group were larger than those from group 2 and 3 (*p* < 0.01 and *p* < 0.05, respectively; **Supplementary Fig. 4 b, d).** Regarding shape, little difference was seen in CAA concavity, however distant cancer adipocyte concavity positively correlated with BMI up to group 3 (*p* ≤ 0.05; **Table 1; Supplementary Fig. 3e, f**). This increase in concavity with BMI was also seen in normal adipocytes from group 2 to 4 (close, *p* < 0.001; distant, *p* ≤ 0.01; **Supplementary Fig. 4e, f**). No differences were seen in aspect ratios, except in breast cancer samples where close adipocytes from the lowest BMI were more circular than those from the highest (*p* ≤ 0.05; **Table 1; Supplementary Fig. 3g, h; Supplementary Fig. 4 g, h**). This analysis shows that adipocyte size but not shape may alter more with BMI, with greater size with increasing BMI more evident in adipocytes from cancer samples than normal samples.

**Table 1.** Size and shape measurements of close adipocytes from the four BMI groups in cancer cases.

| BMI group | Size parameters | Shape parameters |
| --- | --- | --- |

| | Perimeter ($\mu\text{m}$) | Area ($\mu\text{m}^2$) | Concavity ($\mu\text{m}$) | Aspect Ratio |
| --- | --- | --- | --- | --- |
| < 18.5<br>($n = 3$) | 144.91<br>(131.79 – 157.56) | 4868.18<br>(3973.33 – 5670.6) | 369.85<br>(298.588 – 417.35) | 1.32<br>(1.3 – 1.36) |
| 18.5 – 25<br>($n = 17$) | 188.77<br>(165.46 – 212.67)<sup>1</sup> | 6832.2<br>(5219.33 – 8956.94)<sup>1</sup> | 471.28<br>(312.31 – 585) | 1.37<br>(1.33 – 1.42) |
| 25 – 30<br>($n = 10$) | 197.51<br>(173.21 – 238.26)<sup>1</sup> | 8755.8<br>(6638.55 – 10677.6)<sup>1</sup> | 503.1<br>(413.14 – 618.54)<sup>1</sup> | 1.35<br>(1.31 – 1.41) |
| 30 +<br>($n = 20$) | 209.06<br>(174.79 – 247.63)<sup>1</sup> | 7177.25<br>(5750.38 – 9183.7)<sup>1</sup> | 407<br>(292.5 – 573.9) | 1.37<br>(1.32 – 1.43)<sup>1</sup> |

Data expressed as median with interquartile range (IQR). 1 Statistical significance compared to <18.5 (group 1). Full figures, including normal data are in the supplementary material.

### Distant cancer adipocytes are smaller and more elongated in denser breasts

Cancer and normal samples were further assessed to determine whether MD was associated with altered adipocyte morphology. Samples were split according to the BI-RAD (Breast Imaging Reporting and Data) system, which uses the percentage of fibroglandular tissue in relation to adipose tissue to classify breast density into four MD groups: group 1 (< 25% fibroglandular tissue), group 2 (25 – 50% fibroglandular tissue), group 3 (51 – 75% fibroglandular tissue), and group 4 (> 75% fibroglandular tissue) [36]. MD was known for 31 cancer cases: 8 in group 1, 8 in group 2, 9 in group 3, and 6 in group 4. For normal cases, 64 had known MD: 19 in group 1, 16 in group 2, 15 in group 3, and 14 in group 4.

In cancer samples, differences in size were only seen in distant adipocytes, which became smaller with increasing density (perimeter, *p* ≤ 0.01; area, *p* < 0.05; **Table 2; Supplementary Fig. 5a, b, c, d**). This trend was seen in normal samples but only in close adipocytes (*p* ≤ 0.01; **Supplementary Fig. 6a, c**). Distant adipocytes in normal samples had no general trend in size, only that those from group 2 were larger than those from group 3 and 4 (perimeter, *p* ≤ 0.05 and *p* ≤ 0.01, respectively; area, *p* < 0.01; **Supplementary Fig. 6b, d**). No differences were seen in CAA or normal distant adipocyte shape (**Supplementary Fig. 5e, g**; **Supplementary Fig. 6f, h**). In cancer samples, distant adipocytes from the least dense breasts were more concave than those from group 2 and 4 (*p* < 0.001, *p* < 0.01, respectively; **Supplementary Fig. 5f**). Additionally, distant adipocytes from the densest breasts were more elongated than those from the least dense breasts (*p* ≤ 0.05; **Table 2**; **Supplementary Fig. 5h**). In normal samples, close adipocytes from the densest breasts were less concave than those from lower densities (group 1 and 2, *p* < 0.0001; group 4, *p* < 0.001; **Supplementary Fig. 6e**). Adipocyte from group 3 were the most elongated compared with the other groups (group 1 and 4, *p* ≤ 0.01; group 2, *p* ≤ 0.001; **Supplementary Fig. 6g**). These results suggest that changes in MD may be associated with differences in distant adipocyte morphology in cancer samples and close adipocyte morphology in normal samples.

**Table 2.**
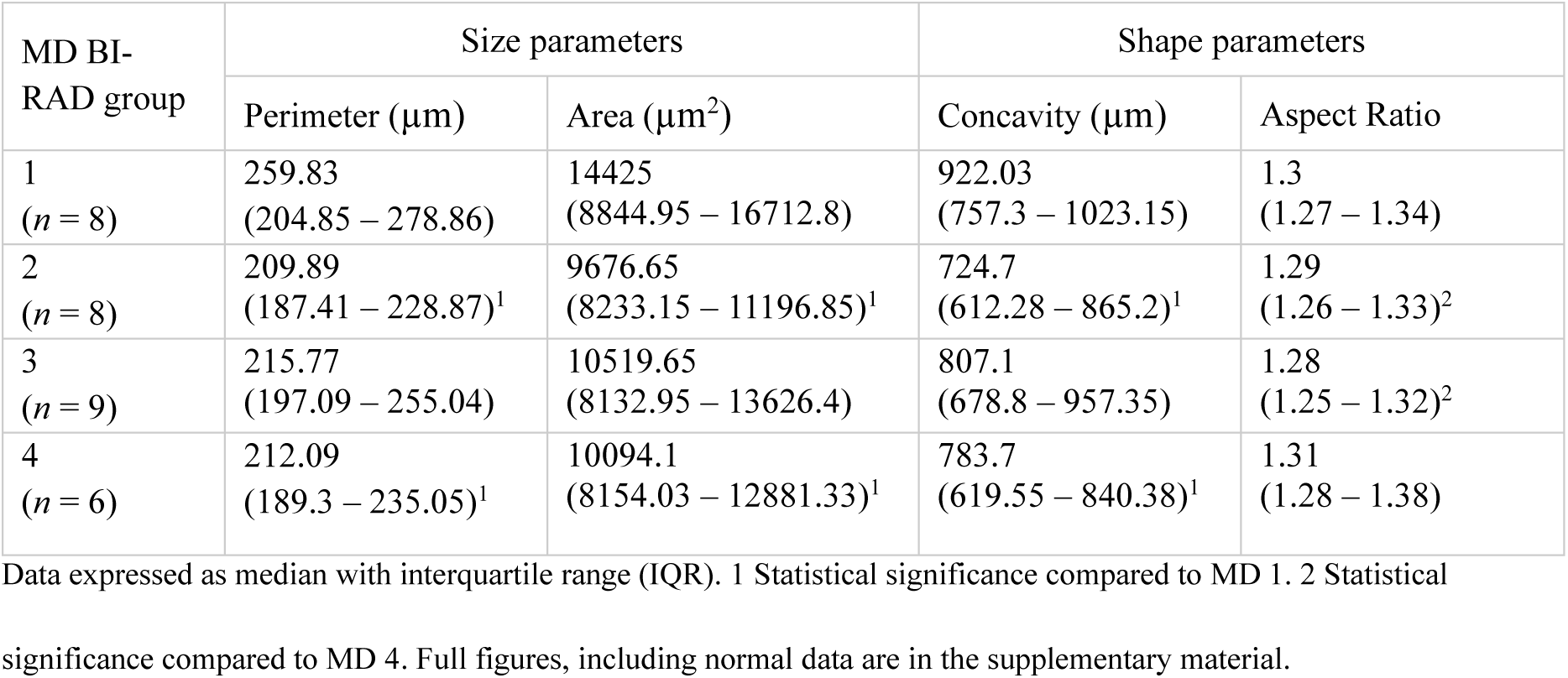
Size and shape measurements of distant adipocytes stratified into the four BI-RAD MD categories in cancer cases.

### Normal adipocytes were larger from donors who later developed breast cancer

Normal, susceptible, breast cancer, and contralateral samples were analysed to determine whether the morphology of normal close adipocytes differed depending on clinical history. Close adipocytes taken from samples from females with no history or diagnosis of breast cancer at the time of sample donation were classed as normal (N; *n* = 116). Those obtained from normal tissue taken from females who later informed Komen Tissue Bank that they had developed breast cancer were classed as susceptible (S; *n* = 29). Adipocytes from the breast with breast cancer were classed as tumour (T; *n* = 123) and those from the breast contralateral to the breast with cancer were classed as contralateral (C; *n* = 29).

Close adipocytes from tumour samples had the smallest perimeter and area compared with those from normal (*p* ≤ 0.0001), susceptible (*p* ≤ 0.0001), and contralateral (perimeter, *p* ≤ 0.001; area *p* ≤ 0.01) samples (**Fig. 3a, b**), as shown in the H&E-stained regions of interest (ROIs) (**Fig. 3e, f, g, h**). Susceptible adipocytes had larger perimeters than normal and tumour adipocytes (*p* ≤ 0.01 and *p* ≤ 0.0001, respectively; **Fig. 3a**), and larger areas than normal, tumour, and contralateral adipocytes (*p* ≤ 0.001, *p* ≤ 0.0001, and *p* ≤ 0.0001, respectively; **Fig. 3b**). Tumour adipocytes were the least concave (*p* < 0.0001, **Fig. 3c**). Interestingly, there was no difference in size between normal and contralateral adipocytes, which were the most circular between the four groups (normal, *p* ≤ 0.01; susceptible, *p* ≤ 0.01; tumour *p* ≤ 0.0001; **Fig. 3d**). These findings indicate that normal close adipocytes were morphologically similar to contralateral adipocytes in size and morphologically distinct from susceptible and tumour adipocytes.

**Fig. 3.**
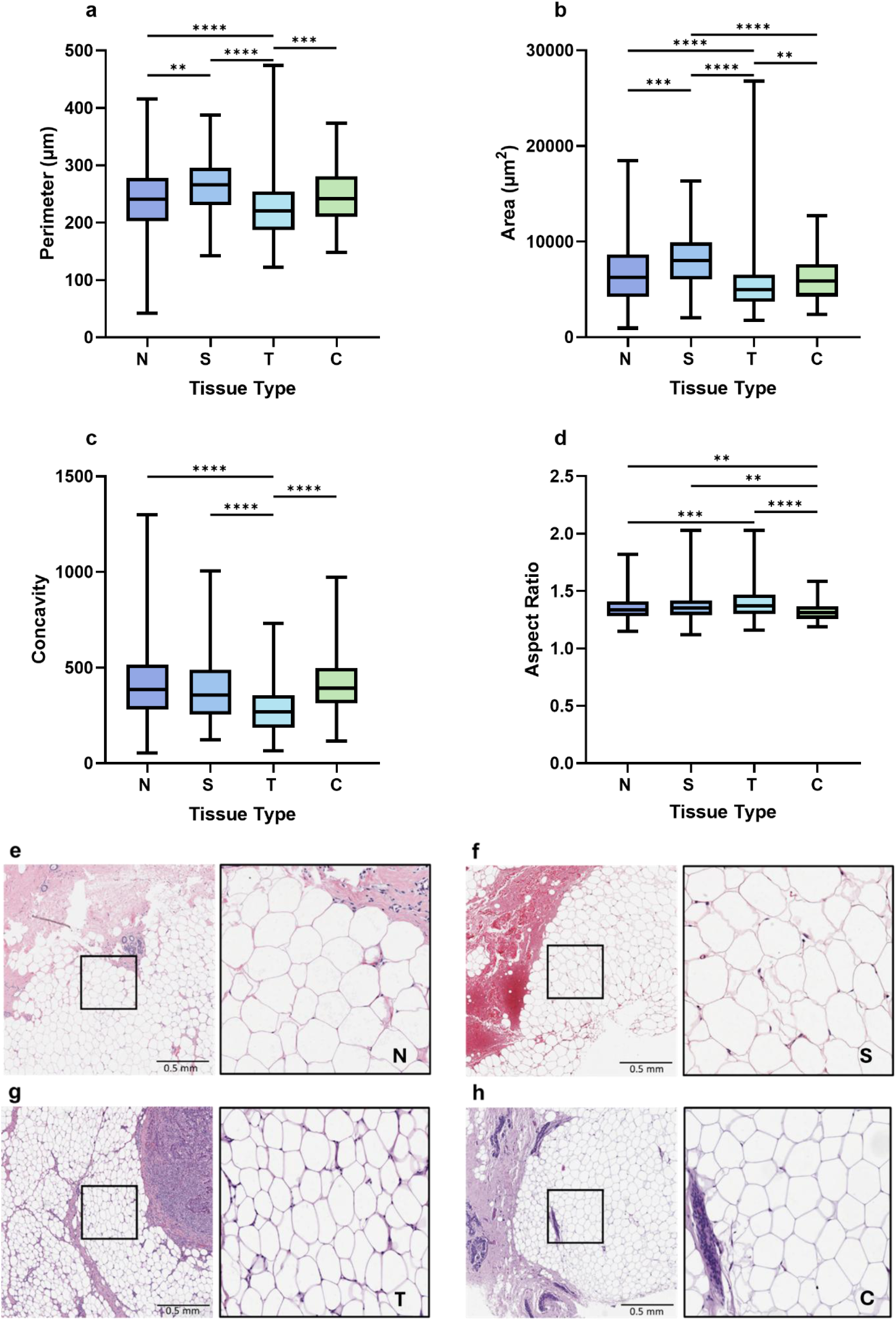
Close adipocytes from normal tissue were morphologically different to adipocytes from susceptible and tumour samples. Median close adipocyte **a** perimeter, **b** area, **c** concavity, **d** aspect ratio from normal (*n* = 116), susceptible (*n* = 29), tumour (*n* = 123), and contralateral (*n* = 29) breast tissue samples. Representative examples of H&E-stained close ROIs from **e** normal, **f** susceptible, **g** breast cancer, and **h** contralateral breast tissue samples. N (normal) = adipocytes from normal breast tissue (*n* = 17,688). S (susceptible) = adipocytes from normal breast tissue from females who later developed breast cancer (*n* = 2,946). T (tumour) = adipocytes from breast cancer tissue (*n* = 17,273). C (contralateral) = adipocytes from the breast contralateral to the breast with cancer (*n* = 8,229). * *p* ≤ 0.05, ** *p* ≤ 0.01, *** *p* ≤ 0.001, **** *p* ≤ 0.0001

### Adipocyte morphology did not impact patient survival

Clinical outcome was known for 68 cancer cases: living (*n* = 50) and deceased (*n* =18). Samples were collected from living donors at the time of tissue collection, with donor outcome collected at the time of analysis. Close adipocytes were split into two groups according to the median values: low (≤ median perimeter = 214.53 µm, area = 4,858.25 µm^2^, concavity = 270.5 µm, and aspect ratio = 1.38) or high (> median values). The same was done for distant adipocytes (median perimeter = 283.82 µm, area = 6051.75 µm^2^, concavity = 468.88 µm, and aspect ratio = 1.28).

Smaller CAAs were associated with poorer survival outcome compared with larger adipocytes when median was used to split the two groups (*p* ≤ 0.05; **Fig. 4a, b**). However, no significant difference was seen in the shape of CAAs and the size and shape of distant adipocytes (**Fig. 4c, d**; **Supplementary Fig. 7**). These results suggest CAAs size may potentially play a role in patient survival, thus additional investigation could provide further insight into whether adipocyte morphology analysis could have clinical applications.

**Fig. 4.**
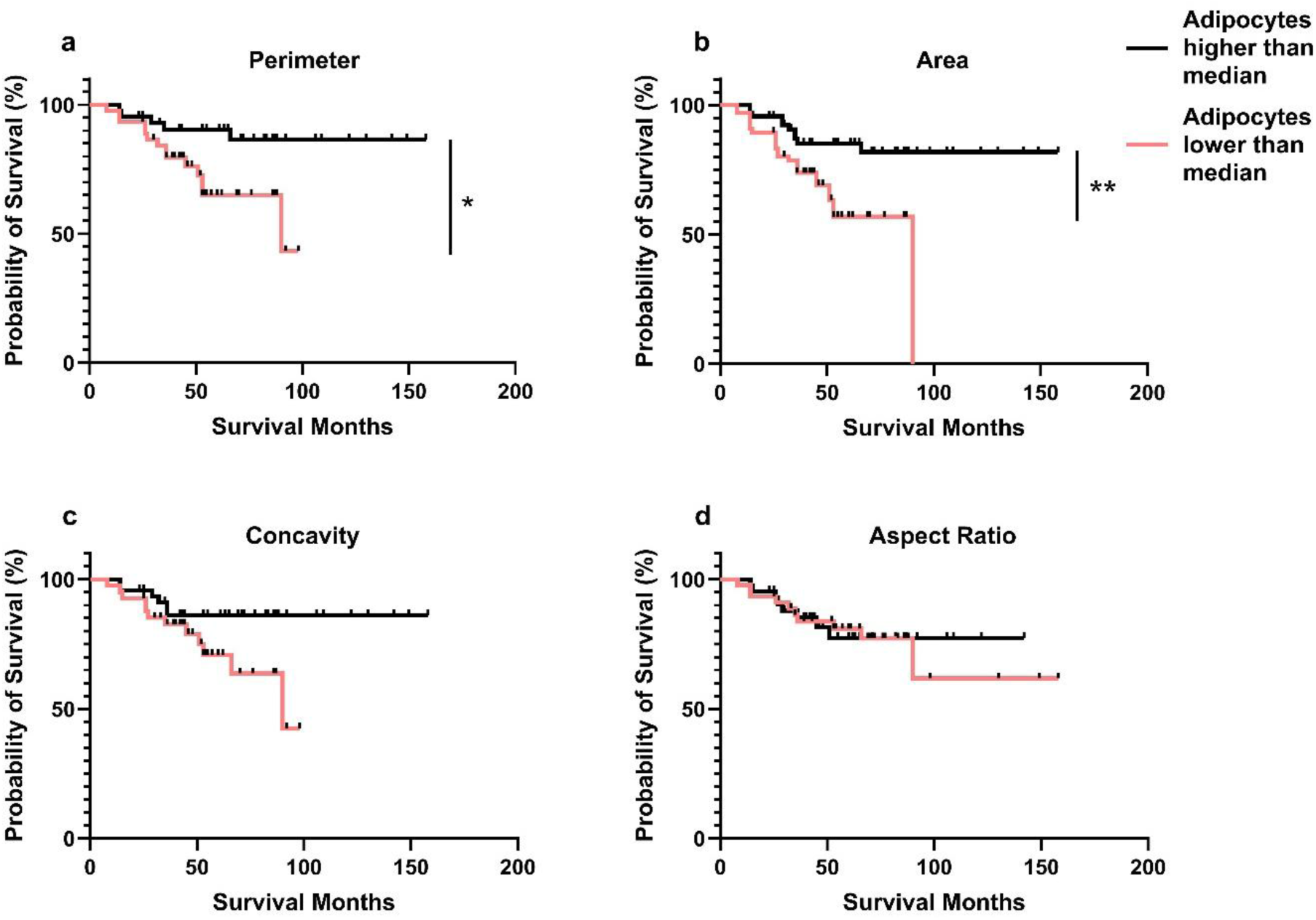
Smaller CAAs were associated with poorer overall survival. Median **a** perimeter, **b** area, **c** concavity, and **d** aspect ratio of close adipocytes from breast cancer tissue samples with known survival outcome (n = 68). Red line = adipocytes lower than median, black line = adipocytes higher than median. Log-rank (Mantal-Cox) test analysis, * *p* ≤ 0.05, ** *p* ≤ 0.01

### Adipocyte morphology differs when stratified into triple-negative and estrogen receptor positive breast cancer subtypes

Adipocyte morphology was assessed in relation to various pathological features, including tumour size, grade and cancer subtype. Tumour size and grade did not impact adipocyte morphology, irrespective of proximity to cancer cells (**Supplementary Fig. 8**). However, when stratified for cancer subtype, adipocytes from triple-negative breast cancer (TNBC; *n* = 36) samples were smaller (*p* < 0.001), more concave (close, *p* < 0.01; distant, *p* < 0.001) and elongated (close, *p* < 0.0001; distant, *p* < 0.05) compared to adipocytes from estrogen receptor positive (ER+; *n* = 22) breast cancer samples (**Supplementary Fig. 9**). This analysis indicates that the tumour size and grade may not influence adipocyte morphology but breast cancer subtype does.

## Discussion

While there has been much progress in defining how cells of the tumour microenvironment communicate with cancer cells, adipocytes have not been considered until much more recently. Adipocyte role in breast cancer has previously been evaluated through the manual measurement of adipocyte diameter using histological tissue sections [18, 19, 37]. However, the size, shape, and contour of adipocytes can be highly irregular in histological sections, thus alternative automated methods have been developed [38–41].

We observed that adipocytes closer to the tumour-leading edge appeared smaller and more fibroblast-like in shape compared with distant adipocytes. This has been reported in other studies, with close adipocytes subsequently being recognised as CAAs [1, 5, 9–11, 42, 43]. Interestingly, this relationship between close and distant adipocytes in breast cancer samples was also seen in normal samples when we evaluated adipocytes based on their proximity to breast epithelial cells. This could be considered the baseline difference between close and distant adipocytes without cancer cell influence. The decrease in close adipocyte size may be a result of collagen stiffness in the extracellular matrix, as reported in a murine study [44]. This suggests that the influence of cancer on CAAs may have been overestimated in previous studies if this baseline was not considered, emphasising the importance of studying normal samples. However, the decrease in size was more than doubled in cancer samples. Given there was no correlation between tumour size and adipocyte morphology, this further decrease may not be caused by tumour compression of adipocytes. Instead, it could be a result of potential bidirectional crosstalk between cancer cells and adipocytes. Breast cancer cells can induce lipolysis within adipocytes to allow for fatty acid transfer and also initiate a loss of terminal differentiation markers in adipocytes [1, 7, 45–48]. This can reduce the volume of the lipid droplet within the adipocytes and thus the overall size, as well as reduce adipocyte circularity as de-differentiation takes place. However, adipocyte morphology changed when various breast cancer risk factors were considered (summarised schematically in **Fig. 5**).

**Fig. 5.**
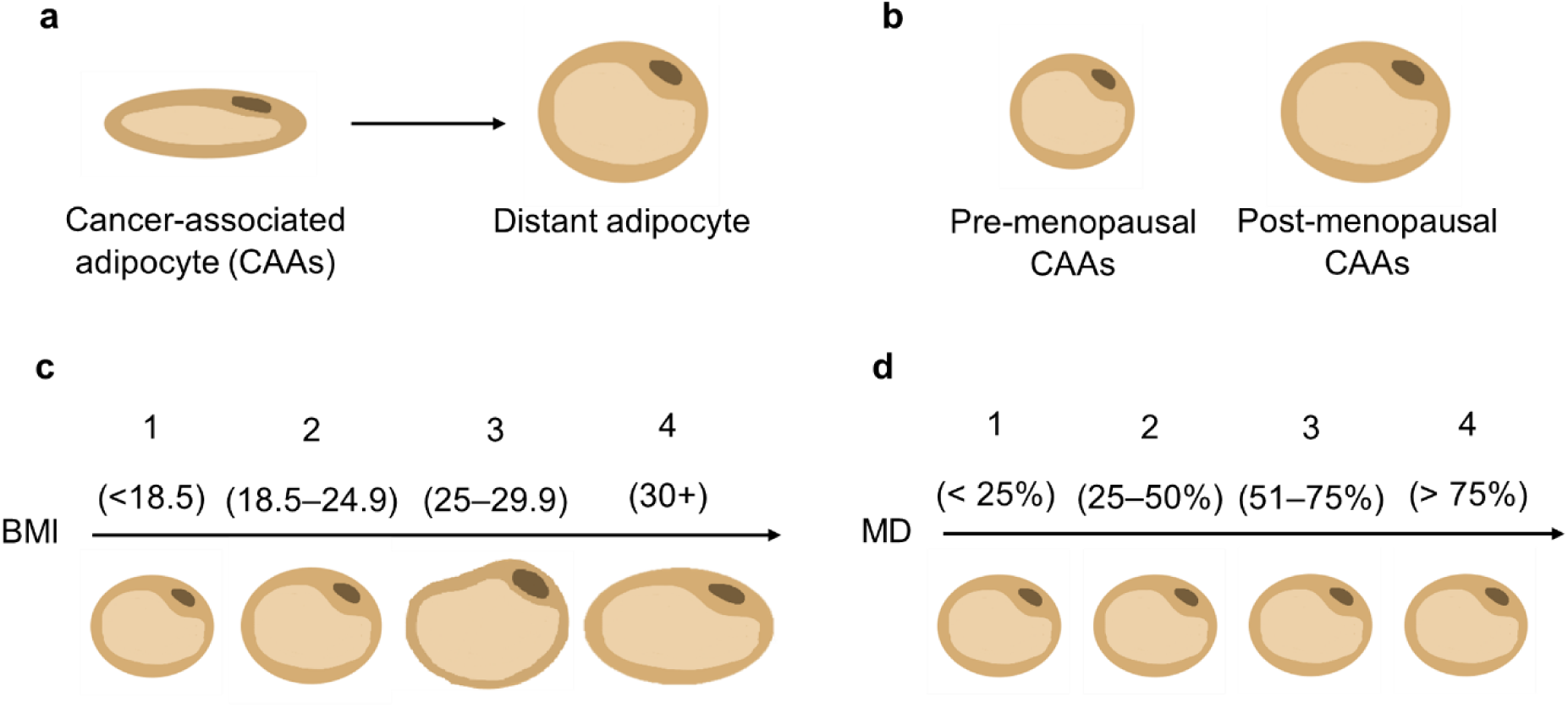
Adipocyte morphology changed depending on the proximity to breast cancer cells as well as the breast cancer risk factor. Adipocyte morphology of **a** CAAs compared with distant adipocytes, **b** pre-menopausal CAAs compared with post-menopausal CAAs, **c** BMI groups 1 - 4 CAAs, and **d** MD groups 1 - 4 CAAs.

When stratifying for menopausal status, CAAs from pre-menopausal cancer samples were smaller than those from post-menopausal samples (**Fig. 5**). However, CAA shape did not differ between the groups. The difference in CAA size might be a result of higher levels of aromatase in post-menopausal females, which is positively correlated with adipocyte diameter [20]. TNBC is more common pre-menopausally, which was reflected in our cohort where 80% of pre-menopausal cases were TNBC compared to 61% of post-menopausal [49]. TNBC is considered more aggressive compared to other breast cancer subtypes [49–52]. Therefore, TNBC could be inducing a higher rate of lipolysis in pre-menopausal females to obtain more energy for growth. This is potentially reflected in our analysis as adipocytes from TNBC samples were smaller and more elongated than those from ER+ samples. Here, we had 20 pre- and 96 post-menopausal cases, with only 5 pre-menopausal samples with known cancer subtypes, making robust sub-group analysis challenging. Differences in lipid composition between pre- and post-menopausal women, along with alterations in cancer cell metabolism, have been linked to shifts in the tumour microenvironment and cancer progression and thus may further contribute to variations in adipocyte morphology [53, 54]. Overall, additional samples should allow for better insights into how menopausal status and cancer subtype may influence adipocyte morphology and thus breast cancer risk and development.

Adipocyte size typically correlates positively with increasing BMI, representing the hypertrophy adipocytes undergo to store excess lipids at a higher BMI (25 +) [15, 55]. Our data showed that this was also seen in breast cancer samples, aligning with previous findings (Fig. 7c) [18]. However, no differences in adipocyte size were observed between the higher BMI groups. It has been reported that normal weight individuals (BMI 18 – 24.9) can be metabolically obese and have larger adipocytes [56]. Additionally, BMI considers muscle and body mass, which is fat-free and can change depending on fitness and age, potentially categorising individuals who are metabolically healthy into the obese group [57–59]. Therefore, this could explain why no differences were seen in adipocyte size between higher BMI groups. Interestingly, when considering shape, adipocytes from the highest BMI were slightly more elongated than those from the lowest. This might result from adipose tissue expansion in obesity caused by the higher lipid uptake in adipocytes, and thus potentially compressing adipocytes into a more elongated shape [60]. Overall, our data supports the notion that BMI might not be an accurate representation of obesity levels. Instead, adipocytes should also be stratified according to other, more accurate measurements, like the waist-hip ratio, to better inform on how obesity may influence adipocyte morphology and to strengthen the validity of our findings.

A high MD is associated with reduced adipose tissue [23, 61]. It could be speculated that having less adipose tissue might result in smaller adipocytes as adipocyte size appeared to decrease in breast cancer tissue as breast density increased [61]. Our analysis supported this in terms of distant adipocytes in cancer samples, but also adipocytes in normal samples, showing that there is an association between MD and adipocyte size irrespective of cancer cell influence. Regarding adipocyte shape, distant cancer adipocytes from the densest breast were more elongated than those from lower densities. The reduced size and elongated shape of adipocytes in denser breasts is potentially due to the higher percentage of fibroglandular tissue in higher MD groups, which may compress adipocytes into becoming smaller and more elongated. Given MD and BMI have an inverse relationship on breast cancer risk and that our results showed both increasing MD and BMI are independently associated with differences in adipocyte size, additional samples are required where both MD and BMI information is available [21–23]. This would allow for further analysis of the joint influence that these two risk factors may have on adipocyte morphology and thus determine the potential importance of incorporating obesity levels into the MD classification system.

Samples of breast tissue from females who later went on to develop cancer (termed susceptible) were used to determine whether adipocyte morphology could infer breast cancer risk. Susceptible breast tissue was previously reported to have a significant increase in lipogenesis and fatty acid transport genes compared with normal tissue [62]. Increased lipogenesis results in increased triglyceride synthesis, leading to a greater lipid uptake by adipocytes. This might explain why, in our study, adipocytes in susceptible breast tissue were larger than those in normal breast tissue obtained from women with no history or diagnosis of breast cancer. Interestingly, these two groups of adipocytes did not differ in shape. However, susceptible adipocytes were more concave than CAAs, suggesting that changes in adipocyte shape may not occur until cancer initiation. It is possible that the differences seen in adipocyte morphology between the various tissue pathologies may be due to patient phenotype as we have shown that menopausal status, BMI, and MD can influence adipocyte morphology. Therefore, matched healthy and susceptible breast tissue would be useful in future experimental studies investigating the underlying reasons for alterations in adipocyte morphology and how adipocytes may play a role in the initiation of breast carcinogenesis.

When adipocyte morphological parameters were categorised into small and large according to the median, smaller CAAs were associated with poorer survival compared with larger CAAs. Bidirectional crosstalk between cancer cells and adipocytes might be an explanation, resulting in the induction of adipocyte lipolysis and transfer of fatty acids to cancer cells [45, 46]. These fatty acids can act as an energy source for cancer cells, possibly enhancing cancer cell growth, and this increase in energy availability may contribute to poorer patient survival [6]. There was no significant difference in survival outcomes depending on the shape of CAAs. For distant adipocytes, neither shape nor size affected survival. However, a recognised limitation was that multivariate analysis was not possible due to incomplete meta data for some cases, including age, BMI, MD, cancer subtype and treatment. Investigation with additional samples could provide insight into the potential of using adipocyte morphology in clinical settings to help assess the risk of disease progression and inform patient outcomes.

In summary, adipocyte morphology was influenced by the location within the breast, the tissue pathology and the various breast cancer risk factors. The use of both cancer and normal samples in this study helped determine the relationship between the risk factors and adipocyte morphology with and without the influence of cancer cells and thus provided a baseline for understanding the extent to which breast cancer cells and associated risk factors impact adipocyte size and shape. In normal samples, adipocyte morphology was influenced more strongly by differences in MD, with close adipocyte size decreasing with increased breast density. However, in breast cancer samples, close adipocyte morphology was more strongly influenced by menopausal status and BMI, with pre-menopausal samples having smaller CAAs compared to post-menopausal samples and CAAs size positively correlating with BMI. Considering the differences in adipocyte morphology, breast cancer risk factors might be influencing the contribution of CAAs to breast carcinogenesis. Overall, this study highlights the importance of studying normal samples and incorporating patient-specific outcomes into investigations on the role of adipocytes in breast cancer development and progression.

## Methods

### Study population

Hematoxylin and eosin (H&E)-stained tissue sections of normal breast tissue samples from females with no history or diagnosis of breast cancer at the point of sample donation, breast cancer, and contralateral breast tissue samples from patients with breast cancer were obtained as whole slide images (WSIs) scanned at x20 magnification from 4 biobanks: Susan G. Komen Tissue Bank (KTB; Subjects were recruited under a protocol approved by the Indiana University Institutional Review Board (IRB protocol number 1011003097 and 1607623663) and according to The Code of Ethics of the World Medical Association (Declaration of Helsinki)), NHS Grampian Biorepository (GB; NHS Grampian Tissue Bank Committee (Network Approval: TR000269)), Breast Cancer Now Biobank (BCNB; BCN 23/EE/0229), and The Leeds Breast Tissue Bank (LBTB; 15/YH/0025), plus open sources WSIs from The Cancer Genome Atlas (TCGA) [63]. From the 537 cases initially identified, 331 were included in the analysis as they met the inclusion criteria shown in **Fig. 6**.

**Fig. 6.**
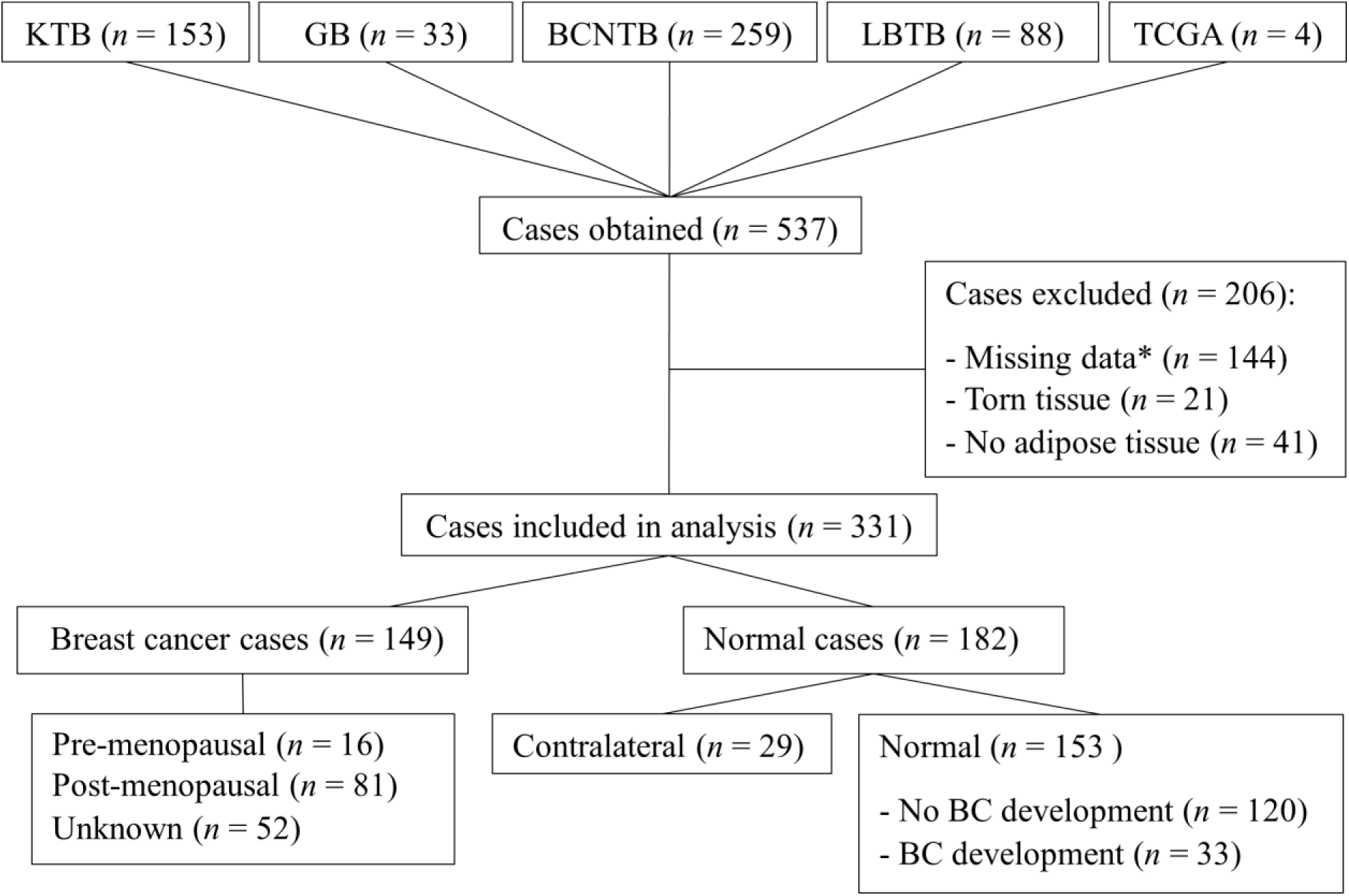
Tissue samples used for adipocyte analysis. A total of 527 digital WSIs of H&E-stained breast cancer, contralateral, and normal breast tissue samples were obtained from five sources. Of these, 331 met the inclusion criteria, with 206 excluded due to missing data as described. Samples were stratified into cancer (*n* = 149) or normal (*n* = 182) groups, and then further split based on menopausal status (pre, *n* = 16; post, *n* = 81; unknown, *n* = 52), tissue pathology (contralateral, *n* = 29; normal, *n* = 153) and breast cancer history (no development, *n* = 120; development, *n* = 33). BC = Breast cancer. * Included age, BMI, MD, and cancer subtype.

### Identification of Regions of Interest and segmentation of adipocytes

Each H&E-stained WSI was uploaded to QuPath open source software [64]. A 500 μm x 500 μm grid was applied and around five ROIs containing close adipocytes (≤ 2 mm away from the tumour-leading edge) and five containing distant adipocytes (>2 mm away) were selected (**Fig. 7a**). The selected areas covered approximately 50 close and 50 distant adipocytes per sample. For the normal breast and contralateral tissue samples, close adipocytes were considered in regions ≤ 2 mm from breast epithelial cells, and distant adipocytes > 2 mm away (**Supplementary Fig. 10**). These distances were chosen to maintain consistency across samples. Similar to cancer samples, five regions were selected of each close and distant adipocytes across the image to ensure the analyses were representative of the whole tissue sample.

ROI images were exported to ImageJ and analysed using the open-source Adiposoft plugin (v1.16) (**Fig. 7b**) [65–68]. Adiposoft automatically segmented adipocytes, excluding those on the edge of the image (**Fig. 7c**). Diameter range of 30 µm - 500 µm was selected to avoid artefacts, such as the lumen of ducts and blood vessels. The background was cleared and adipocyte annotations were filled before the image was converted to an 8-bit binary image, a pre-requisite for the ImageJ Particles8 plugin (**Fig. 7d, e**) [69].

**Fig. 7.**
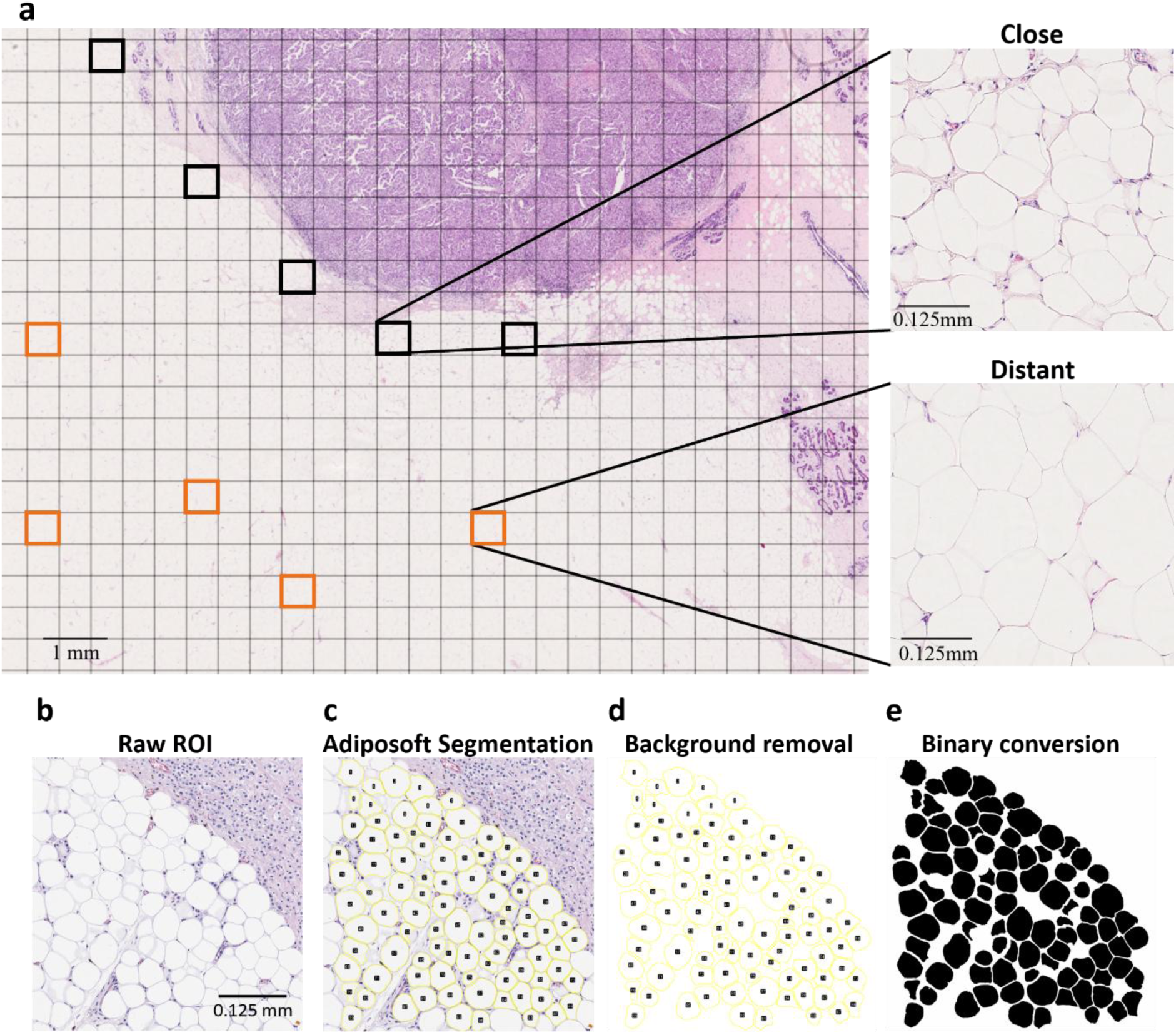
Image analysis performed on WSI of H&E-stained breast cancer tissue sample and CAAs ROI. **a** Five ROIs were selected of close adipocytes (black squares) ≤ 2 mm from the tumour-leading edge and of distant adipocytes (orange squares) > 2 mm away. **b** TIFF image of H&E-stained ROI of close adipocytes from a WSI of a H&E-stained breast cancer tissue sample was loaded into ImageJ and the Adiposoft plugin was applied. Adipocytes on the edge of the image were excluded and only those of a diameter between 30 – 500 µm were included. Following **c** automatic segmentation, **d** the image background was removed, **e** and the adipocyte outlines filled. The image was converted to an 8-bit binary image. WSI scanned at x20 magnification.

### Analysis of morphological features

Adiposoft segmentation generated 51,751 CAAs, 27,003 distant cancer adipocytes, 7,746 close and 2,798 distant contralateral adipocytes, and 21,318 close and 11,274 distant normal adipocytes. Particles8 plugin was used to analyse the binary images. Based on previous literature, parameters that were the most biologically informative were calculated from each adipocyte [70]. Out of 7 parameters, perimeter, area, concavity, and aspect ratio were further assessed as these were the most descriptive parameters for representing adipocyte morphology. Measurements were converted from pixels to microns using the scale bar. The median for each parameter was calculated from each ROI.

### Statistical analysis and identification of important parameters

A Random Forest analysis was conducted (R Statistical Software v4.4.0; R Core Team 2024) to determine which of the 7 biologically informative parameters from the Particles8 analysis were the most important for evaluating adipocyte morphology (**Supplementary Fig. 11**) [71]. Literature suggested that area and diameter were the most common descriptors of size [55, 60, 72, 73]. Diameter, given as Feret by Particles8, was lower than area and perimeter in terms of accuracy and reliability of the model (**Supplementary Fig. 11a, b**). Given perimeter was the highest size parameter in the model, it was used to describe adipocyte size along with area for comparison to existing literature. Concavity, which shows how concave the adipocytes are, was the most important shape parameter and thus was used within this study (**Supplementary Fig. 11a, b**). Removal of area, perimeter, or concavity meant the overall accuracy and reliability of the model in classifying the adipocytes as either close or distant notably decreased. Even though aspect ratio was less important within the model, it was used in this study as it is calculated using two of the 7 biologically informative parameters (feret and breadth) and gives insight into whether adipocytes were circular or more elongated like CAAs [5, 45]. Definitions of the four parameters (perimeter, area, concavity, and aspect ratio) used in this study are provided in **Table 3**.

**Table 3.**
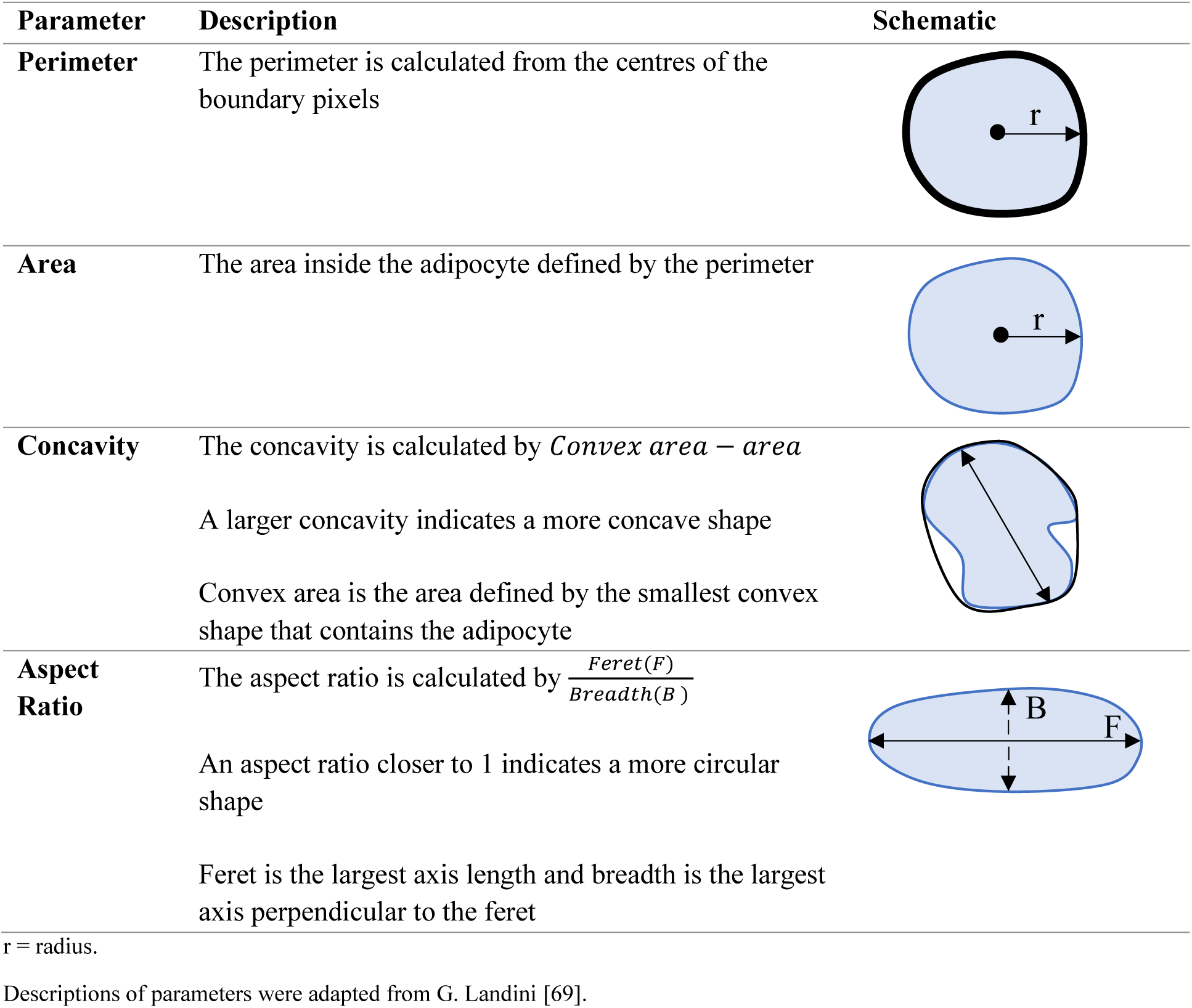
Description and schematic representation of the most discriminant Particles8 parameters evaluated.

| Parameter | Description | Schematic |
| --- | --- | --- |
| **Perimeter** | The perimeter is calculated from the centres of the boundary pixels |  |
| **Area** | The area inside the adipocyte defined by the perimeter |  |
| **Concavity** | The concavity is calculated by $Convex\ area - area$<br>A larger concavity indicates a more concave shape<br>Convex area is the area defined by the smallest convex shape that contains the adipocyte |  |
| **Aspect Ratio** | The aspect ratio is calculated by $\frac{Feret(F)}{Breadth(B)}$<br>An aspect ratio closer to 1 indicates a more circular shape<br>Feret is the largest axis length and breadth is the largest axis perpendicular to the feret |  |

r = radius.
Descriptions of parameters were adapted from G. Landini [69].

Shapiro Wilk normality test followed by Kruskal-Wallis H with Dunn’s multiple comparison tests were used to determine statistical differences in adipocyte morphology. *P* values ≤ 0.05 were considered statistically significant. Statistical tests and graphs were done using Graph Pad Prism version 10.0.3. for Windows.

## Ethics Approval

Ethics approval was covered for the four biobanks: Susan G. Komen Tissue Bank, NHS Grampian Biorepository, Breast Cancer Now Biobank, and The Leeds Breast Tissue Bank. Ethics approval was not required from the open source The Cancer Genome Atlas (TCGA) as images obtained from TCGA were from tissues collected from fully anonymised patients in a publicly available dataset.

## Competing Interests

The authors report there are no competing interests to declare.

## Acknowledgements

We thank Breast Cancer Now Biobank, The Leeds Breast Tissue Bank, The Grampian Biorepository, and Susan G. Komen Tissue Bank at the Indiana University Simon Cancer Center for providing access to H&E stained WSIs, as well as donors and their families, whose help and participation made this work possible. Special thanks to The Cyril and Margaret Gates Charitable Trust for contributing to this work. The results published here are in part based upon data generated by the TCGA Research Network: https://www.cancer.gov/tcga.

## Funding

AD, KH and KP were funded by The Cancer PhD Programme at The University of Aberdeen, an Elphinstone Scholarship, and Animal Free Research UK/Breast Cancer UK co-fund, respectively.

## Contributions

VS, RAE., and JR. conceived the study. AD, KH, KP, and IH acquired and analysed data and prepared figures. GL, NS, EH, and BE provided tissues and images. AD, KH, KP, RAE, JR, and VS drafted the manuscript. All authors reviewed and contributed to the final manuscript.

## Data Availability

Data generated or analysed during this study is available from the DOI: https://doi.org/10.6084/m9.figshare.29040689

## Notes

https://doi.org/10.6084/m9.figshare.29040689

### Competing Interest Statement

The authors have declared no competing interest.

