## Supplementary File for "A compendium of adipocyte morphologies across different breast pathologies"

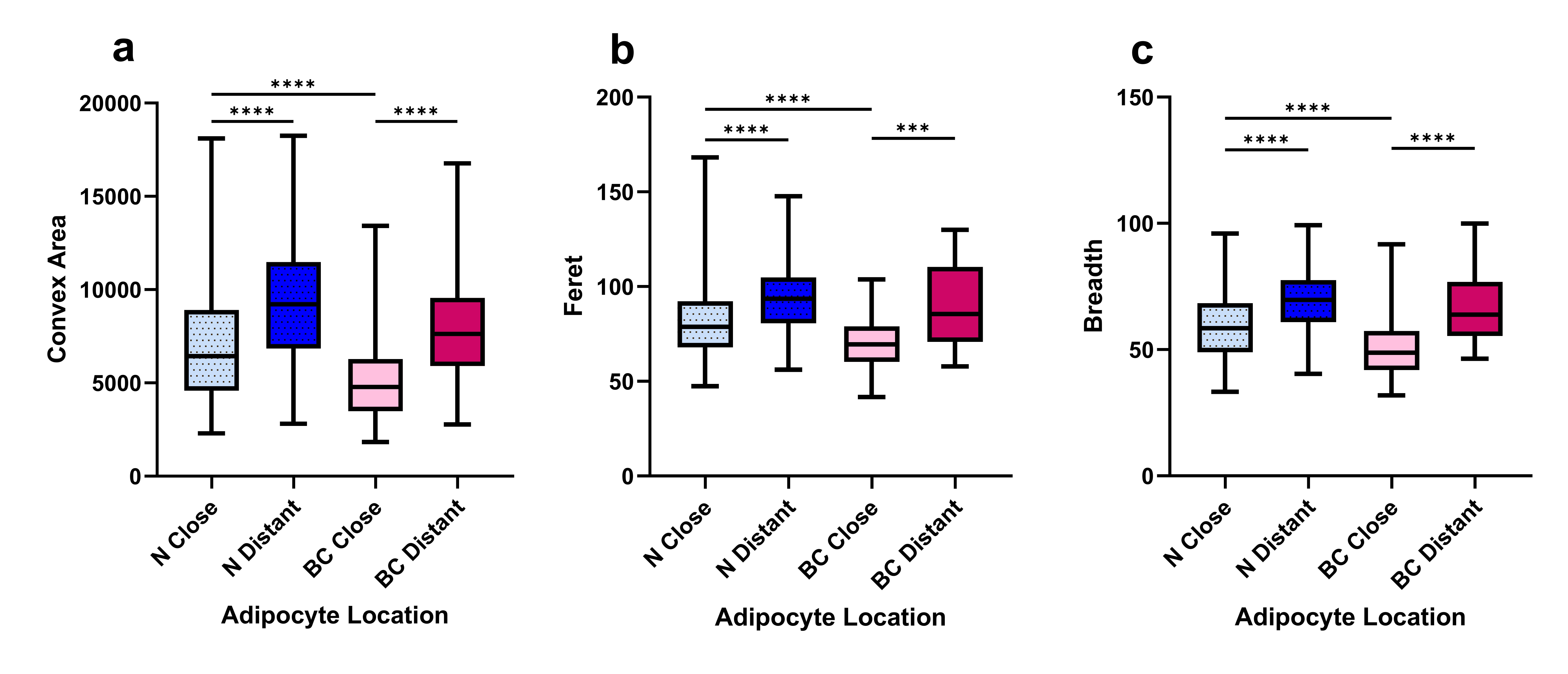

**Supplementary Fig. 1** Additional parameters confirmed findings that CAAs were smaller than normal close adipocytes. Median adipocyte **a** convex area, **b** feret, and **c** breadth from breast cancer (*n* = 149) and normal (*n* = 153) samples. N Close = normal adipocytes ≤ 2 mm away from the breast epithelial cells (*n* = 21,318). N Distant = normal adipocytes > 2 mm away from the breast epithelial cells (*n* = 11,274). BC Close = breast cancer adipocytes ≤ 2 mm away from the tumour-leading edge (*n* = 51,751). BC Distant = breast cancer adipocytes > 2 mm away from the tumour-leading edge (*n* = 27,003). *** *p* ≤ 0.001, **** *p* ≤ 0.0001

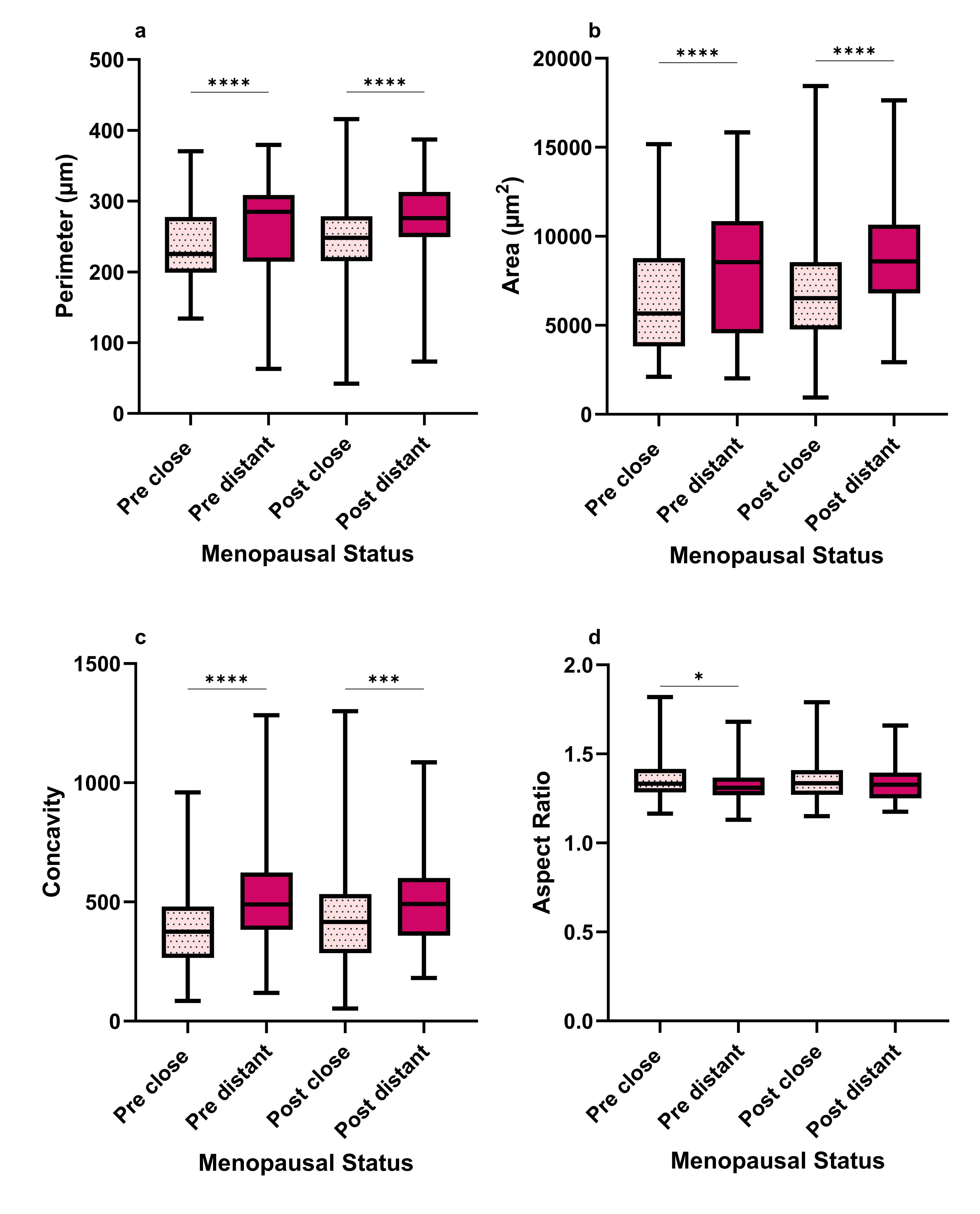

**Supplementary Fig. 2** Size and shape did not differ between pre- and post-menopausal close adipocytes in normal breast tissue. Median adipocyte **a** perimeter, **b** area, **c** concavity, and **d** aspect ratio from normal breast tissue samples (*n* = 130). Pre = Pre-menopausal samples (*n* = 48). Post = Post-menopausal samples (*n* = 82). Close = adipocytes ≤ 2 mm away from the breast epithelial cells (pre, *n* = 7,876; post, *n* = 9,877). Distant = adipocytes > 2 mm away from the breast epithelial cells (Pre, *n* = 3,512; post, *n* = 5,743). * *p* ≤ 0.05, *** *p* ≤ 0.001, **** *p* ≤ 0.0001

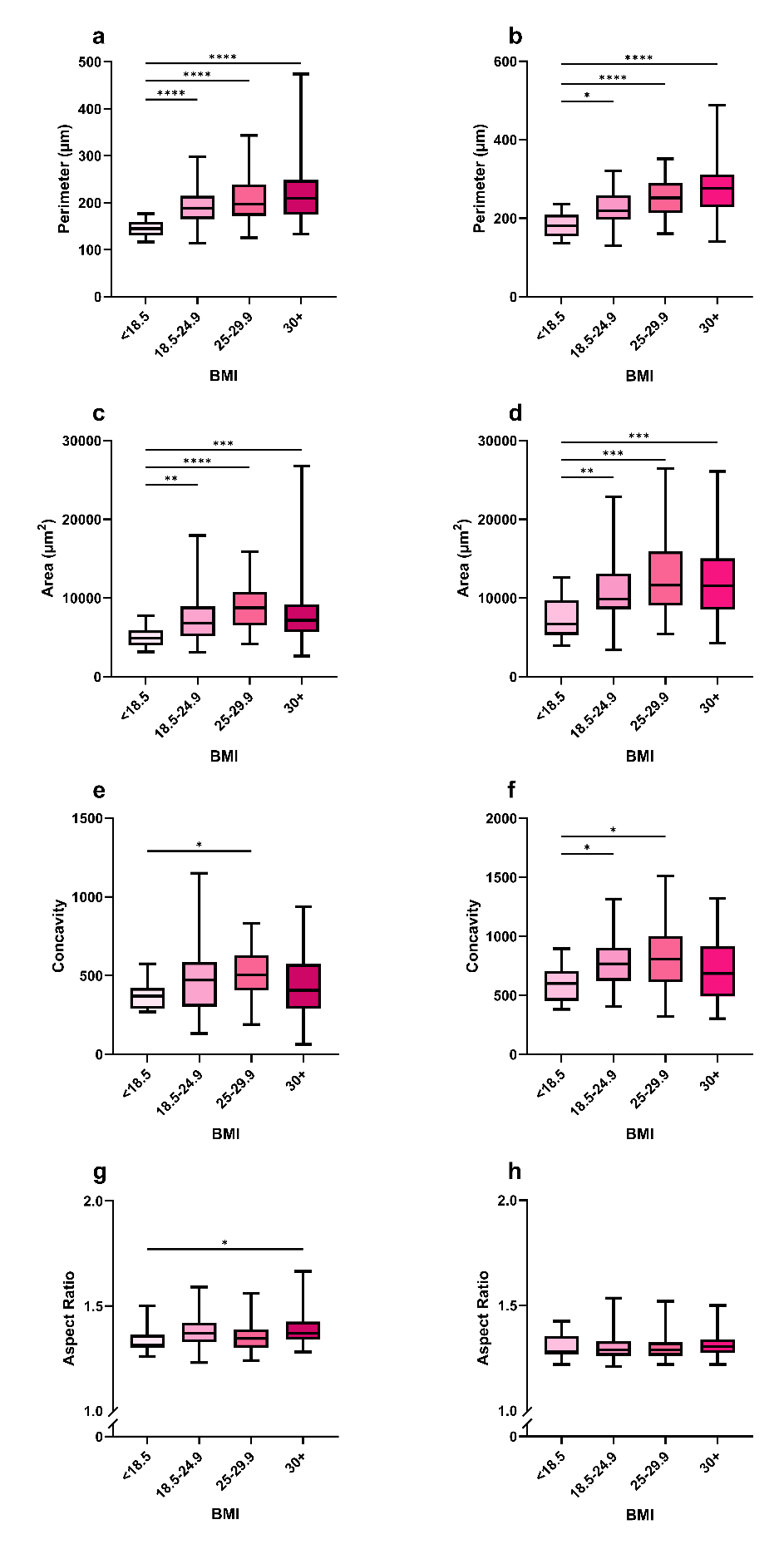
**Supplementary Fig. 3** Close and distant adipocytes increased with BMI in breast cancer samples. Median **a, b** perimeter, **c, d** area, **e, f** concavity, and **g, h** aspect ratio of **a, c, e, g** close (< 18.5, *n* = 4,445; 18.5 – 24.9, *n* = 13,192; 25 – 29.9, *n* = 6,938; 30 +, *n* = 11,251) and **b, d, f, h** distant (< 18.5, *n* = 3,058; 18.5 – 24.9, *n* =10,242; 25 – 29.9, *n* = 5,319; 30 +, *n* =6,025) adipocytes from breast cancer tissue samples (n = 50). * *p* 0.05, ** *p* ≤ 0.01, *** *p* ≤ 0.001, **** *p* ≤ 0.0001

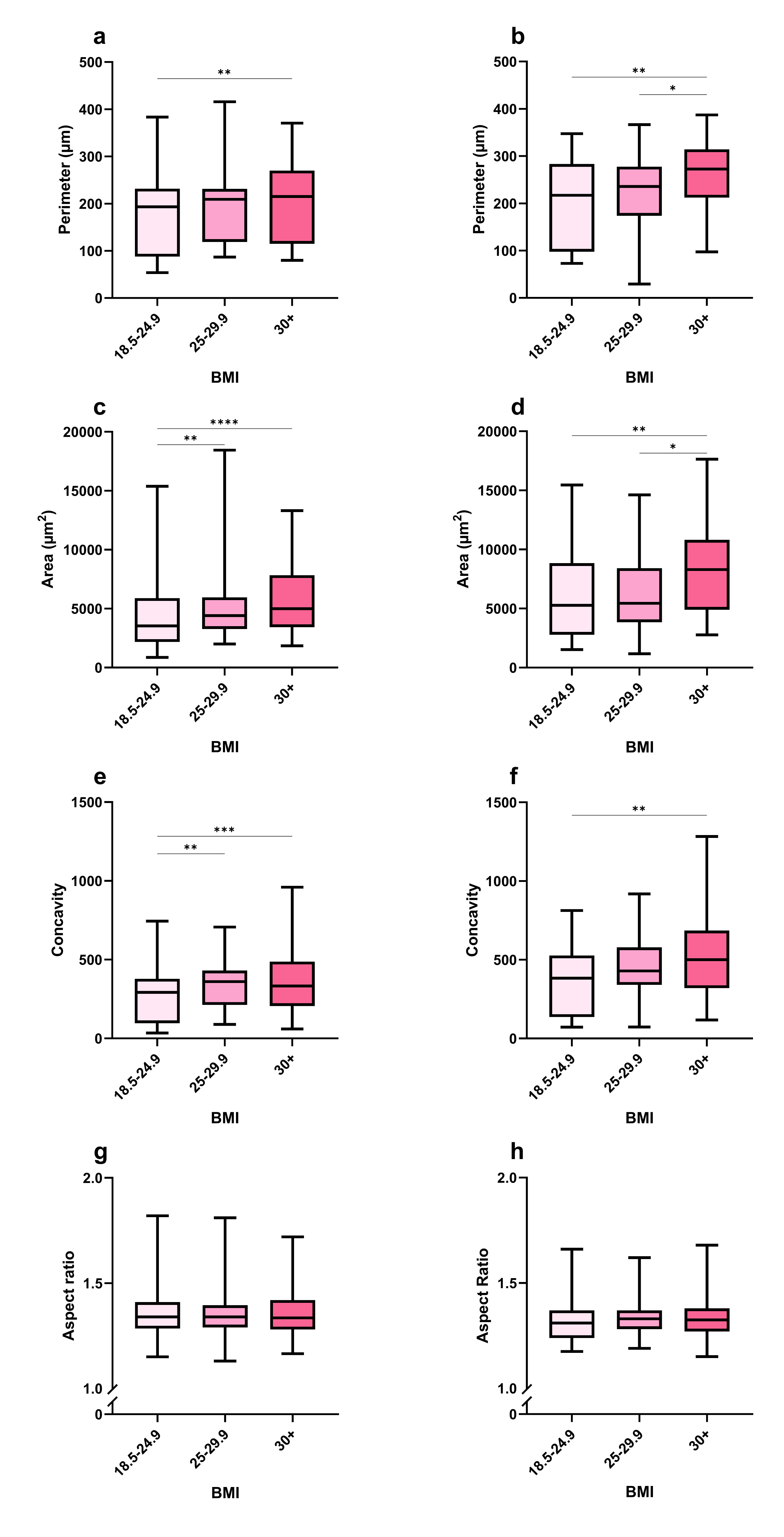
**Supplementary Fig. 4** Limited differences were seen in normal adipocyte morphology when stratifying for BMI. Median **a, b** perimeter, **c, d** area, **e, f** concavity, and **g, h** aspect ratio of **a, c, e, g** close (18.5 – 24.9, *n* = 3,915; 25 – 29.9, *n* = 2,302; 30 +, *n* = 1,991) and **b, d, f, h** distant (18.5 – 24.9, *n* = 1,507; 25 – 29.9, *n* = 2,346; 30 +, *n* = 3,262) adipocytes from normal tissue samples (n = 84). * *p* 0.05, ** *p* ≤ 0.01, *** *p* ≤ 0.001, **** *p* ≤ 0.0001

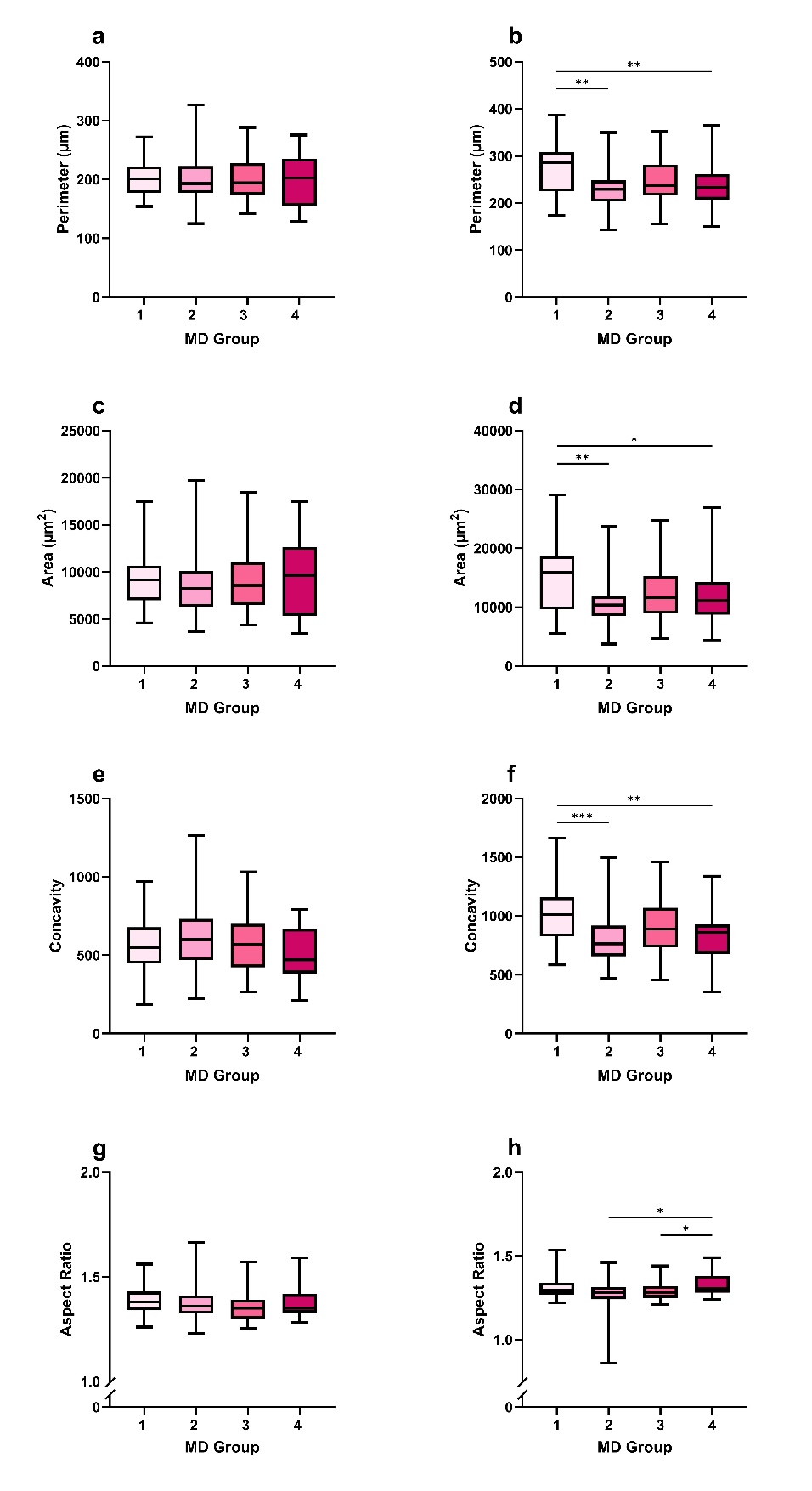

**Supplementary Fig. 5** Distant adipocyte size increased with MD in breast cancer samples. Median **a, b** perimeter, **c, d** area, **e, f** concavity, and **g, h** aspect ratio of **a, c, e, g** close (group 1, *n* = 5,978 ; group 2, *n* = 7,325; group 3, *n*  = 10,460; group 4, *n*  = 6,943) and **b, d, f, h** distant (group 1, *n* = 4,670; group 2, *n* = 6,411; group 3, *n*  = 6,906; group 4, *n*  = 5,088) adipocytes from breast cancer samples (*n* = 31). MD groups were determined using the BI-RAD system. * *p* ≤ 0.05, ** *p* ≤ 0.01, *** *p* ≤ 0.001, **** *p* ≤ 0.0001

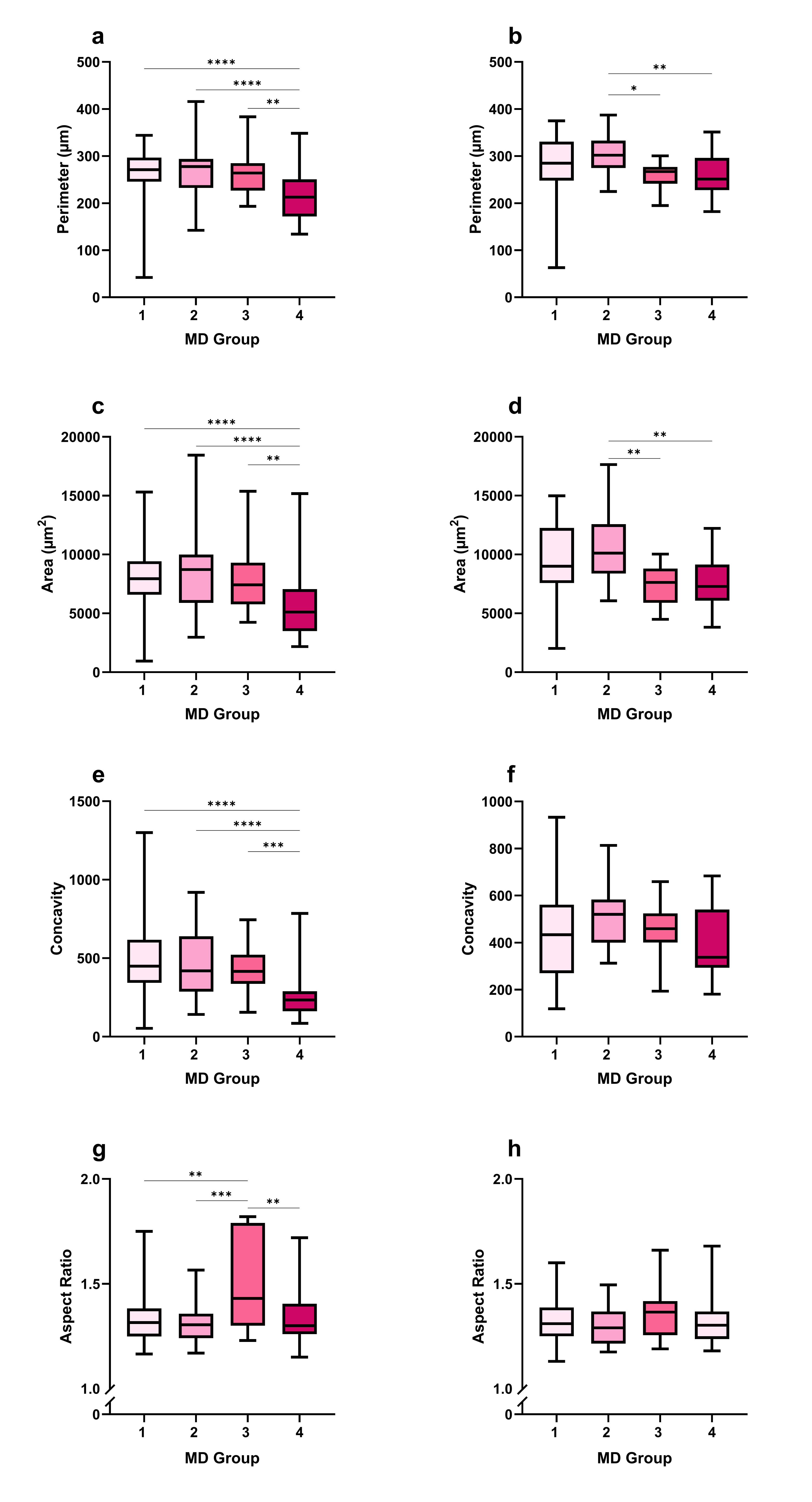

**Supplementary Fig. 6** Close adipocyte morphology differed more between MD groups compared to distant adipocyte morphology in normal samples. Median **a, b** perimeter, **c, d** area, **e, f** concavity, and **g, h** aspect ratio of **a, c, e, g** close (group 1, *n* = 1,877; group 2, *n* =998; group 3, *n*  = 736; group 4, *n* = 2,586) and **b, d, f, h** distant (group 1, *n* = 750; group 2, *n* = 499; group 3, *n*  = 445; group 4, *n* = 745) adipocytes from normal breast samples (*n* = 64). MD groups were determined using the BI-RAD system. ** *p* ≤ 0.01, *** *p* ≤ 0.001, **** *p* ≤ 0.0001

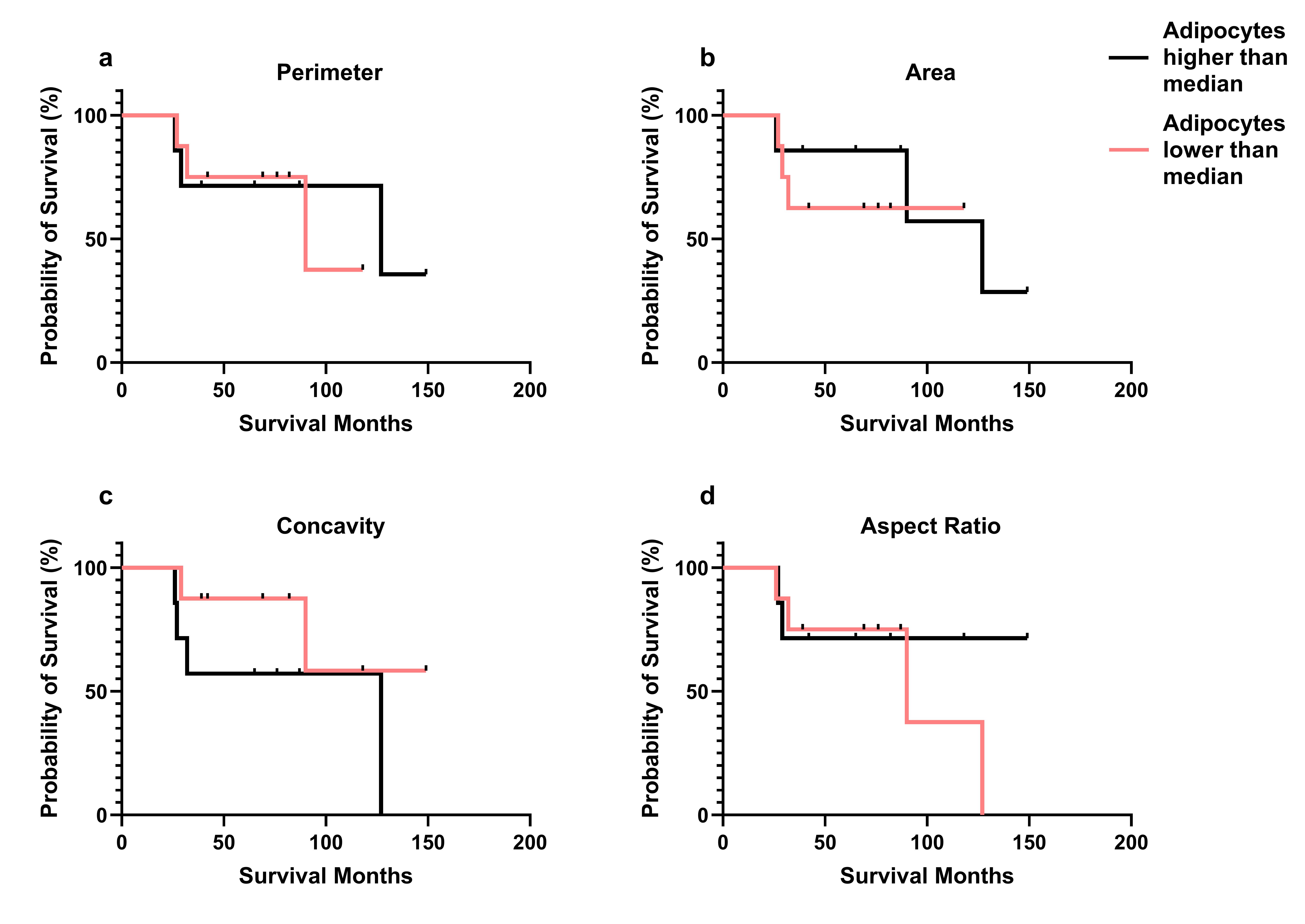
**Supplementary Fig. 7** Distant adipocyte morphology was not associated with overall survival. Median **a** perimeter, **b** area, **c** concavity, and **d** aspect ratio of distant adipocytes from breast cancer tissue samples with known survival outcome (n = 68). Red line = adipocytes lower than median, black line = adipocytes higher than median.

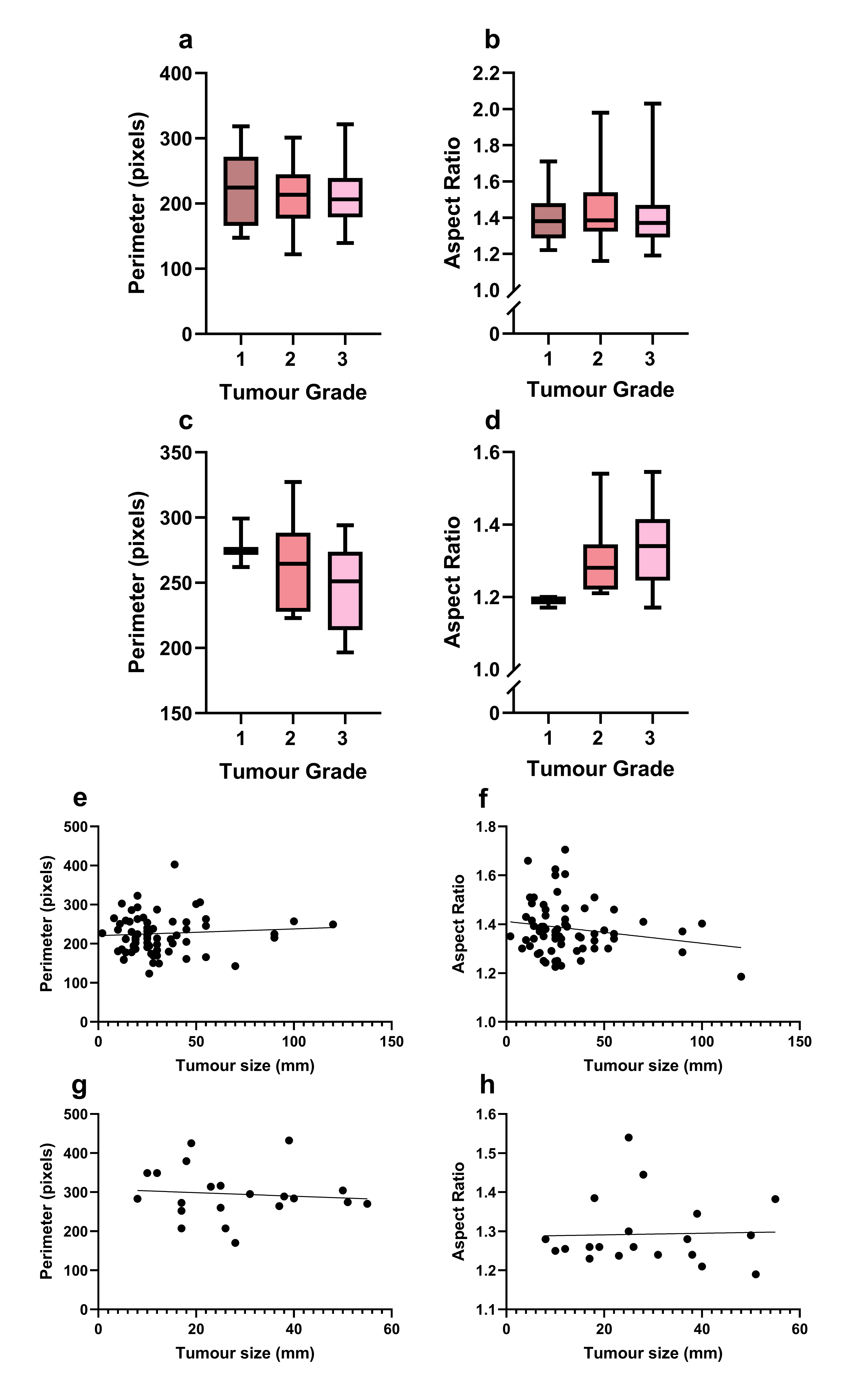

**Supplementary Fig. 8** Tumour grade and size did not influence adipocyte morphology irrespective of location within the breast. Adipocyte **a, c, e, g** perimeter and **b, d, f, h** aspect ratio of **a, b, e, f** close and **c, d, g, h** distant adipocytes from breast cancer samples with known **a – d** tumour grade (grade 1, *n*  = 10; grade 2, *n*  = 16, grade 3, *n* = 38) and **e – h** tumour size (*n* = 70).

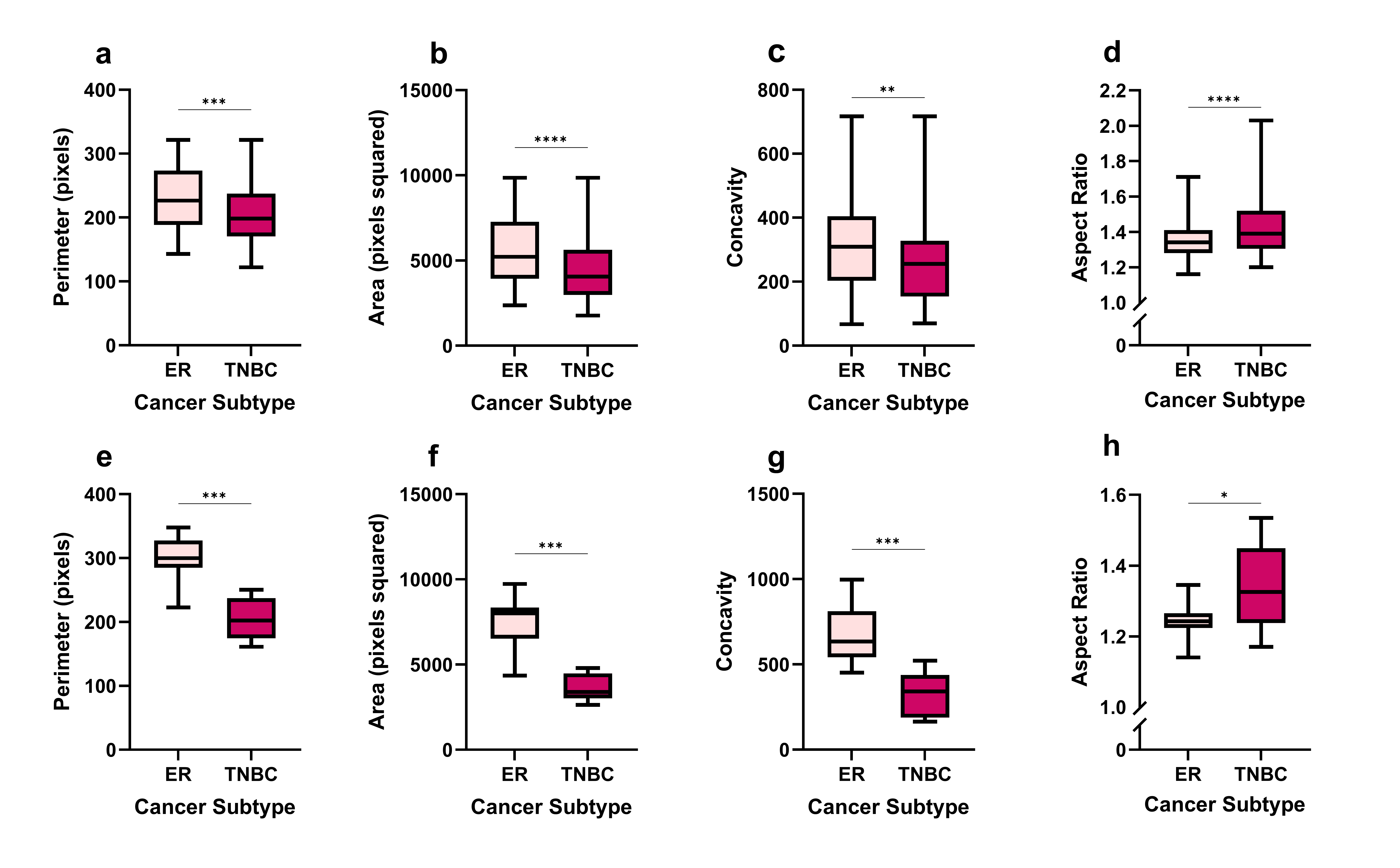

**Supplementary Fig. 9** Adipocytes were smaller and more elongated from TNBC breast cancer tissue samples compared to ER+ samples. Median **a, e** perimeter, **b, f** area, **c, g** concavity, and **d, h** aspect ratio from **a – d** close and **e – f** distant adipocytes from breast cancer tissue samples with known cancer subtypes (ER+, *n* = 22; TNBC, *n* = 36).

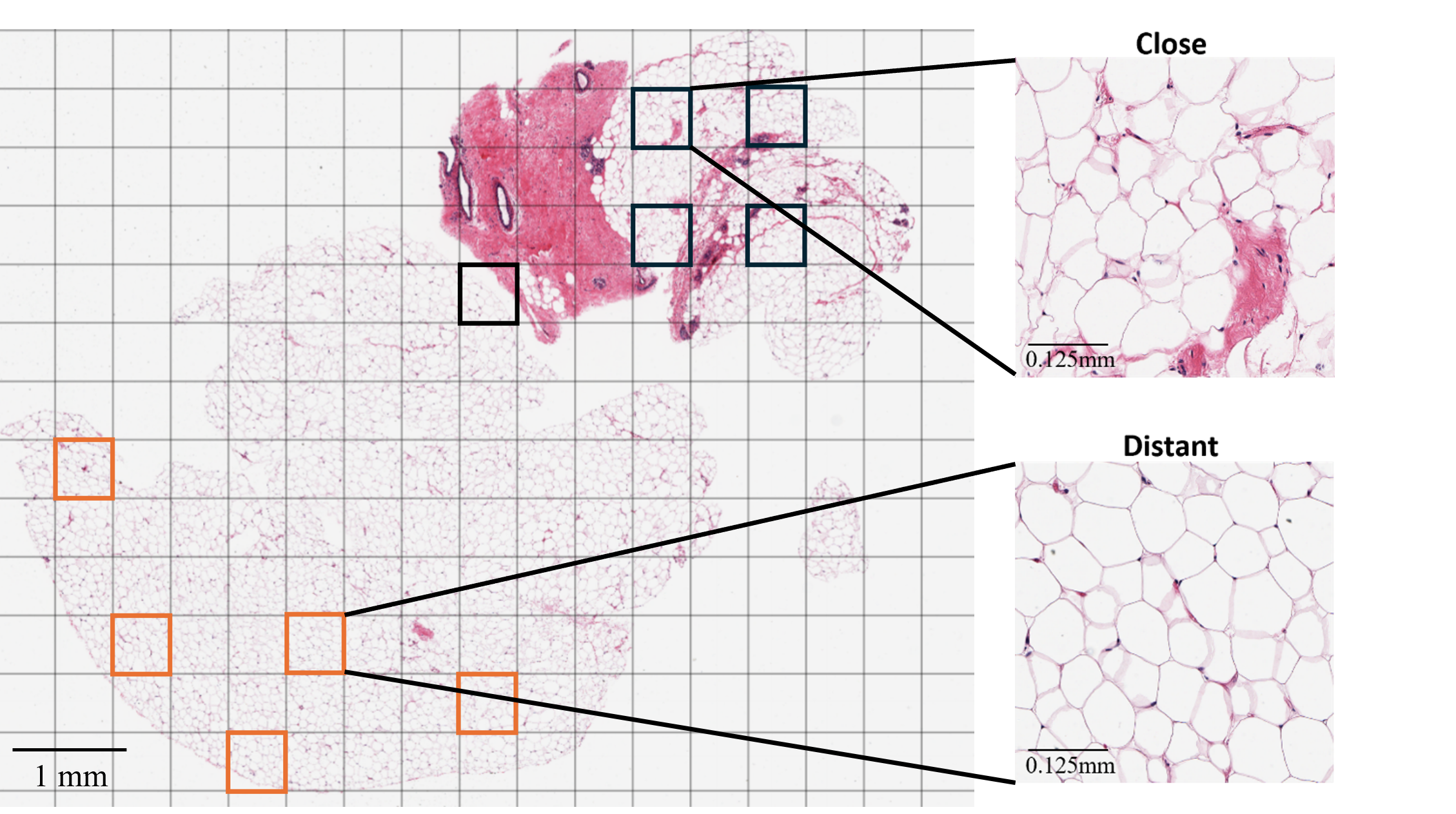

**Supplementary Fig. 10** WSI of a H&E-stained normal breast tissue sample with close and distant ROI. Five ROI were selected of close adipocytes (black squares) ≤ 2 mm from the breast epithelial cells and of distant adipocytes (orange squares) > 2 mm away. Image scanned at x20 magnification.

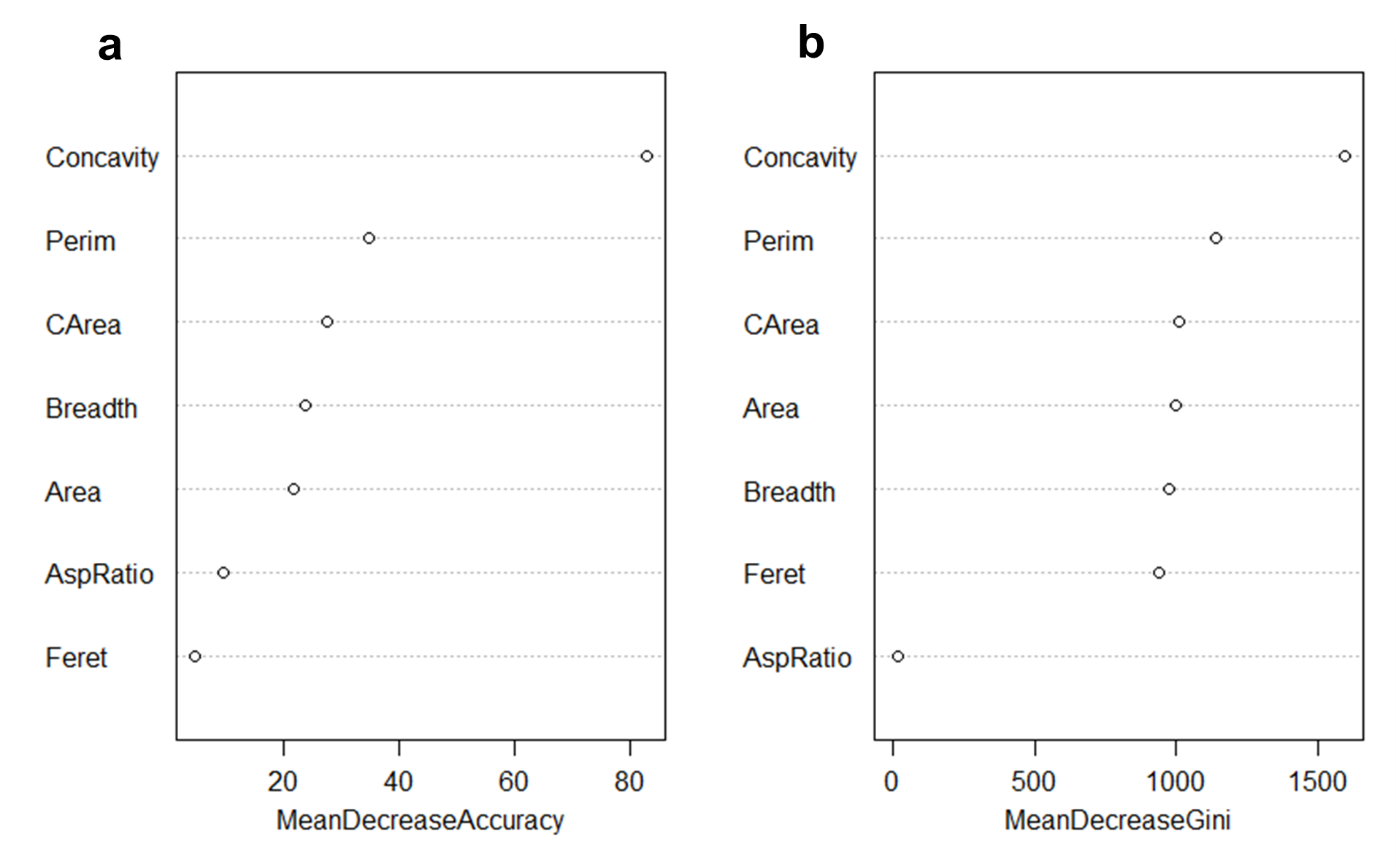
**Supplementary Fig. 11** Random Forest analysis showing the most important Particles8 parameters in classifying adipocytes as close or distant to breast cancer cells. **a** MeanDecreaseAccuracy shows which parameter would decrease the accuracy of the Random Forest model in splitting adipocytes into close or distant the most when removed. **b** MeanDecreaseGini indicates which parameter contributes the most to making the Random Forest model reliable at separating close and distant adipocytes. Analysis conducted using R Statistical Software (v4.4.0; R Core Team 2024).
